# Evolution of miRNA binding sites and regulatory networks in cichlids

**DOI:** 10.1101/2021.12.14.472604

**Authors:** Tarang K. Mehta, Luca Penso-Dolfin, Will Nash, Sushmita Roy, Federica Di-Palma, Wilfried Haerty

## Abstract

The divergence of regulatory regions and gene regulatory network (GRN) rewiring is a key driver of cichlid phenotypic diversity. However, the contribution of miRNA binding site turnover has yet to be linked to GRN evolution across cichlids. Here, we extend our previous studies by analysing the selective constraints driving evolution of miRNA and transcription factor (TF) binding sites of target genes, to infer instances of cichlid GRN rewiring associated with regulatory binding site turnover. Comparative analyses identified increased species-specific networks that are functionally associated to traits of cichlid phenotypic diversity. The evolutionary rewiring is associated with differential models of miRNA and TF binding site turnover, driven by a high proportion of fast-evolving polymorphic sites in adaptive trait genes compared to subsets of random genes. Positive selection acting upon discrete mutations in these regulatory regions is likely to be an important mechanism in rewiring GRNs in rapidly radiating cichlids. Regulatory variants of functionally associated miRNA and TF binding sites of visual opsin genes differentially segregate according to phylogeny and ecology of Lake Malawi species, identifying both rewired e.g. clade-specific and conserved network motifs of adaptive trait associated GRNs. Our approach revealed several novel candidate regulators, regulatory regions and three-node motifs across cichlid genomes with previously reported associations to known adaptive evolutionary traits.

## Introduction

The molecular ‘tinkering’ of ancestral systems and divergence of gene regulatory processes is a hallmark of evolution, and has long been thought to be associated with morphological diversity (Wilson et al. 1974; King and Wilson 1975; Prager and Wilson 1975; Jacob 1977). Based on these theories, a number of studies have focused on gene regulatory networks (GRNs) with the aim of relating gene expression variation to phenotypic divergence (Carroll 2000; Carroll 2008; Peter and Davidson 2011). With this aim, we recently developed an integrative approach to comparatively study GRN evolution across multiple tissues along a phylogeny (Mehta et al. 2021). However, our previous approach largely focused on gene co-expression and transcription factor binding site (TFBS) evolution, without assessing the contribution of other regulatory mechanisms towards GRN evolution, like posttranscriptional repression. This process generally occurs at the three prime untranslated region (3’ UTR) of a gene, which can contain binding sites for both RNA-binding proteins (RBPs) and small non-coding RNAs (ncRNAs), such as microRNAs (miRNAs). miRNAs are key regulators of gene expression, and therefore fundamental to the evolution of novel phenotypes across the animal kingdom (Berezikov 2011).

Vertebrate clades differ dramatically in species richness, and ray-finned fishes represent the largest radiation of any group (>32,000 species). Amongst this radiation, the East African cichlids are a diverse clade that arguably represents the most speciose example of adaptive radiations. In the three great lakes of East Africa (Tanganyika, Victoria and Malawi) and within the last 10 million years (Genner et al. 2007; Wagner et al. 2012), one or a few ancestral lineages of cichlid fish have independently radiated into over 2000 species. These species have been able to explore a variety of ecological niches and partly as a result (Wagner et al. 2012), have given rise to an explosive diversity of phenotypic traits (Kocher 2004). Using genome and transcriptome sequences of five representative East African species, we previously demonstrated that a number of molecular mechanisms may have contributed to diversification, including the rapid evolution of regulatory elements and the emergence of novel miRNAs that may alter gene expression programmes (Brawand et al. 2014). Recent studies, focused on genomic analysis of a wider range of lake species, identified low levels (0.1-0.25%) of genetic diversity between Lake Malawi species pairs (Malinsky et al. 2018), and link species richness in Lake Tanganyika tribes to variable heterozygosity, but not to the accelerated evolution of coding sequences (Ronco et al. 2021). Investigations of Lake Victoria species have also highlighted the role of ancient indel polymorphisms in non-coding regions towards species ecological divergence (McGee et al. 2020). These findings largely report that the genomes are very similar within same lake species. This implies that discrete differences, like regulatory changes, are likely to have an important role in controlling gene expression and function, ultimately contributing to the large phenotypic differences among species. Indeed, our comparative approach focused on the integration of gene co-expression and TFBS motifs in promoter regions, to characterise GRN evolution in six tissues of five East African cichlids (Mehta et al. 2021). We identified GRN changes along the phylogeny, including cases of network rewiring for visual genes (Mehta et al. 2021). We experimentally validated that TFBS mutations have disrupted regulatory edges across species, and segregate according to lake species phylogeny and ecology (Mehta et al. 2021). These findings suggested that GRN rewiring could be a key contributor to cichlid phenotypic diversity (Mehta et al. 2021).

By using similar techniques to those applied to study transcription factors (TFs) (Thompson et al. 2015), previous studies in cichlids have only focused on mRNA/miRNA expression and sequence evolution at miRNA binding sites. Previous analyses reported signatures of purifying selection on cichlid miRNA binding sites (Franchini et al. 2016; Kautt et al. 2020), and that on average, cichlid 3’ UTRs were longer with more miRNA targets per gene than in non-cichlid teleost species (Xiong et al. 2018). Conserved miRNAs tend to differ across species in their expression levels, sequence, distribution and number of predicted binding sites (Xiong et al. 2019). Additionally, there is also evidence for the acquisition of between 36 and 1738 novel miRNAs in the rapidly radiating cichlids (Brawand et al. 2014; Franchini et al. 2016; Franchini et al. 2019; Xiong et al. 2019) and for a higher evolutionary rate of 3’ UTR divergence among cichlid species (Xiong et al. 2018). Genes of the longest and most rapidly evolving 3’ UTRs were found to be associated with translation and ribosomal pathways (Xiong et al. 2018).

No previous studies have analysed the selective constraint of miRNAs and their targets in cichlids. This can be assessed by studying the turnover of miRNA binding sites, which can be defined as the rate at which an ancestrally conserved miRNAs acquire novel binding sites or lose existing ones along a phylogeny. Previous studies in other organisms identified more targets for older, than younger miRNAs in *Drosophila* (Nozawa et al. 2016), conserved regulatory roles for conserved miRNAs in primates (Simkin et al. 2014), and three characteristic rates of target site gain and loss during mammalian evolution (Simkin et al. 2020).

Despite the role of miRNAs as key gene regulators, the dynamic turnover of their binding sites in vertebrates (Simkin et al. 2014; Simkin et al. 2020), and the potential role of GRN rewiring as a key contributor to East African cichlid phenotypic diversity (Mehta et al. 2021), no previous study has explored the contribution of miRNAs and miRNA binding site turnover towards GRN rewiring events across cichlids. Instead, our previous work characterised a single layer (transcriptional activation) of cichlid GRNs solely based on gene co-expression data and predicted gene promoter TFBS interactions (Mehta et al. 2021). In this study, we use our previously published genomic datasets (Brawand et al. 2014) and predicted GRNs in five East African cichlids (Mehta et al. 2021), with the aims of 1) extending the cichlid GRNs with an additional layer (post-transcriptional repression) based on predicted miRNA-mRNA interactions; 2) integrate and analyse nucleotide conservation and/or variation at miRNA binding sites to better understand the selective constraints driving their evolution; 3) characterise co-regulation of target genes by TFs and miRNAs as three-node motifs to study wider GRN evolution; 4) infer instances of three-node motif and GRN rewiring attributed to regulatory binding site turnover; and 5) analyse the plausibility of whether TF and miRNA binding site turnover could be associated with traits of cichlid phenotypic diversity.

## Results

### miRNA binding site prediction in 3’ UTRs of genes

Building upon our previously characterised cichlid GRNs based on TFBSs (Mehta et al. 2021), and to specifically assess miRNA binding site turnover on GRN rewiring and ultimately contributions towards cichlid phenotypic diversity, we used 992 cichlid miRNA mature sequences from 172 families (Brawand et al. 2014) to predict miRNA binding sites in five cichlid species using Targetscan7 (Agarwal et al. 2015). To predict high-confidence miRNA targets, we used the Targetscan7 context++ model for miRNA targeting efficacy (Agarwal et al. 2015). Using a weighted context++ score threshold of < −0.1 (see *Materials and Methods*), like that previously applied in other studies of vertebrate miRNA binding sites (Warnefors et al. 2017; Hu et al. 2019; Sayed and Park 2020), we predicted 19,613,903 miRNA binding sites in the 3’ UTRs of 21,871 orthogroups across five cichlid species (see *Materials and Methods*, Fig. 1a). We further filtered our data to only include 3’ UTRs from 18,799 co-expressed orthogroups to match our previous data set for downstream analyses (Mehta et al. 2021), resulting in a total of 15,390,993 predicted binding sites across the five species (Fig. 1a, Supplementary Fig. S1). Using these predicted binding sites, we classified unique predicted miRNA binding sites of a target gene (TG) 3’ UTR as a miRNA:TG edge in each species, and compared the total number of common and unique miRNA binding sites across all orthologous TGs based on miRNA:TG overlap (Fig. 1b, see *Supplementary information*). We note that there are 33,814 common sites between all species and that the three haplochromine species share the second most number (16,164) of binding sites (Fig. 1b). Unbiased by genome completeness or annotation quality (see *Supplementary information)*, between 31,186 (*P. nyererei*) and 128,831 (*A. burtoni*) unique binding sites were found to be unique to a species (Fig. 1b). In total, 3’ UTR binding sites are predicted for 172 miRNA families (*M. zebra* – 118; *P. nyererei* – 117; *A. burtoni* – 151; *N. brichardi* – 115; and *O. niloticus* – 129). For instance, *miR-15c* binding sites are under-represented in *N. brichardi* (Supplementary Fig. S2). This could be attributed to mutations of the *miR-15c* seed sequence in *N. brichardi* (AGCAGCG) as compared to the other species (AGCAGCA) (see *Supplementary information*). Gene ontology (GO) enrichment of target genes for the miRNA families highlight terms that are both common e.g. membrane and signal transduction, and unique e.g. ATP binding and zinc ion binding (FDR<0.05) between the five species (Supplementary Fig. S3). Overall, variation in the number of binding sites and GO enrichment of the 172 miRNA families across the five species supports differential targeting of genes in each species.

**Fig. 1.**
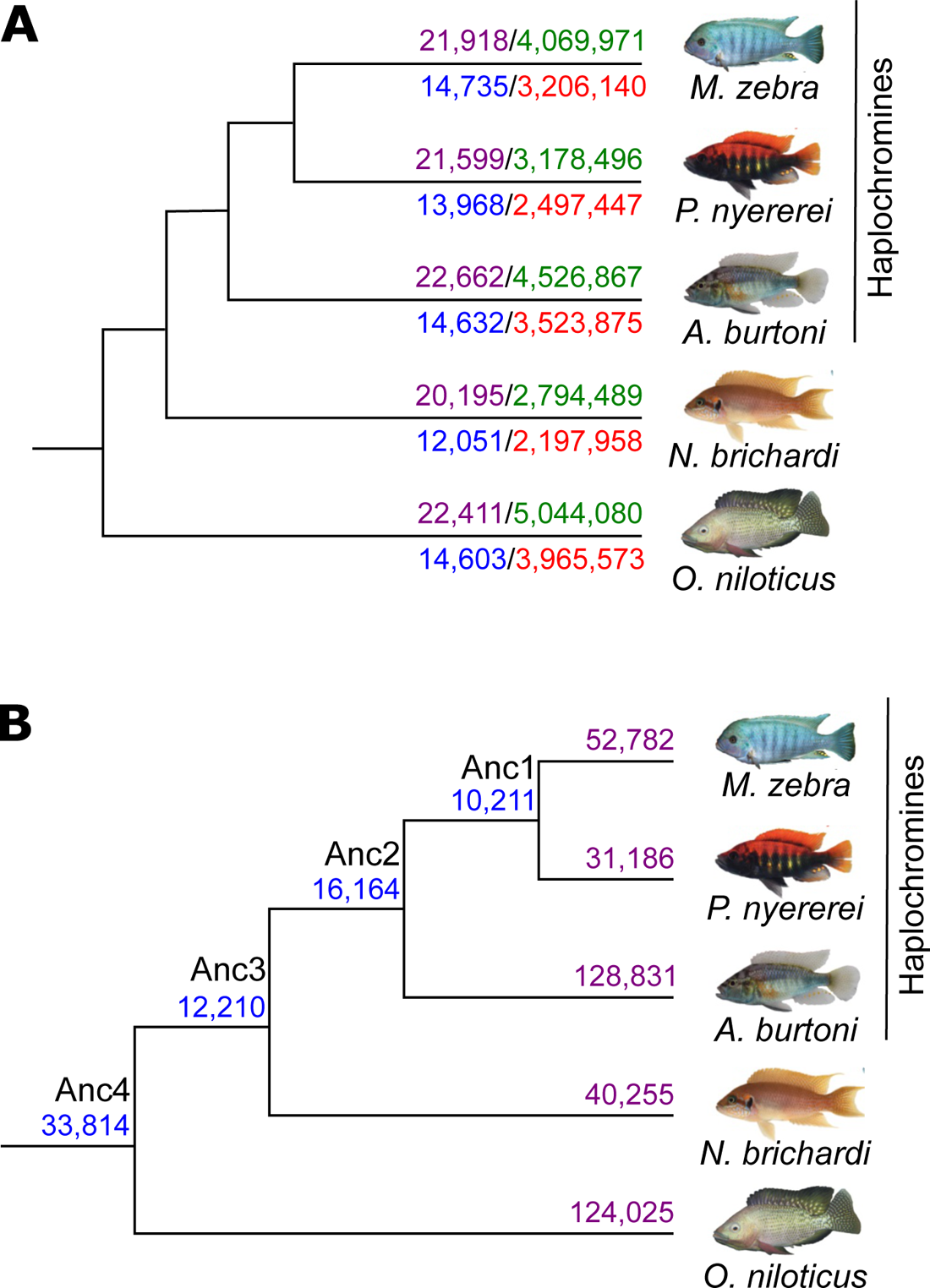
miRNA target prediction in five cichlid species. **(A)** Number of miRNA binding sites predicted across 3’ UTR sequences in each species. Number of all input orthogroup 3’ UTR sequences for each species (purple numbers) and predicted miRNA binding sites from TargetScan7 after filtering for low quality predictions (green numbers) are shown for each species above the branch. Number of 3’ UTR sequences across 18,799 co-expressed orthogroups for each species (blue numbers) and predicted miRNA binding sites (red numbers) are shown for each species below the branch. **(B)** Number of common and unique miRNA binding sites across 3’ UTR sequences of co-expressed orthogroups. Number of common miRNA target sites across 3’ UTR sequences of 18,799 co-expressed orthogroups are shown at ancestral nodes (blue numbers) and unique binding sites in each species (purple numbers). Common and unique binding sites at each node are defined based on overlap of unique miRNA family and target gene edges between species (see *Supplementary information*).

### Differential miRNA binding site usage highlights rewiring at the post-transcriptional level

To study miRNA binding site usage, we assess binding site conservation and divergence based on overlap of aligned 3’ UTR regions (Supplementary Fig. S4). If at the same or overlapping positions in the alignment, a binding site has been predicted for more than one miRNA family between at least two species, then the ancestral binding site is predicted to be functionally diverged (see *Supplementary information* and Supplementary Fig. S4). Compared to an average nucleotide identity of 95 – 99.7% across coding sequences, representative of genomic regions under strong selective pressure, the average nucleotide identity across all 3’ UTR alignments ranges from 83 - 95% across all pairwise species comparisons. This is similar to the average nucleotide identity of 85 – 89% across pairwise comparisons of each species whole genome, representative of the average selective pressure across all genomic regions. By filtering targets based on complete positional overlap in at least two species, we retained a total of 1,626,489 3’ UTR binding sites across all species (18,626/18,799 orthogroups represented). To predict functional divergence, we classified unique predicted miRNA binding sites of a target gene (TG) 3’ UTR as a miRNA:TG edge in each species, and assessed the number of shared sites (in orthologous TGs) utilised by miRNA families that are either the same (miRNA:TG overlap - Fig. 2a, Supplementary Fig. S5a) or different (no miRNA but only TG overlap - Fig. 2b, Supplementary Fig. S5b) between species. Consistent with the previous findings (Fig. 1b), most sites (50,212) are conserved across all species (Anc4 node, Fig. 2a). Following the phylogenetic relationships, the haplochromine species share the second highest number (Anc2 node: 32,087) of binding sites (Fig. 2a). Overall, binding sites are generally conserved and utilised by orthologous miRNA families along the whole phylogeny. Counter to this, compared to basal phylogenetic comparisons (Anc4:1 and Anc3:17 shared sites), there is more miRNA family divergence within the haplochromine lineage (Anc2:3163 and Anc1:3200 shared sites) (Fig. 2b). For example, the developmental gene, *gata6*, has one miRNA binding site (*miR-27d*) shared between *N. brichardi* and *O. niloticus*, but in the haplochromines, has three miRNA binding sites (*miR-219, miR-128* and *miR-27*).

**Fig. 2.**
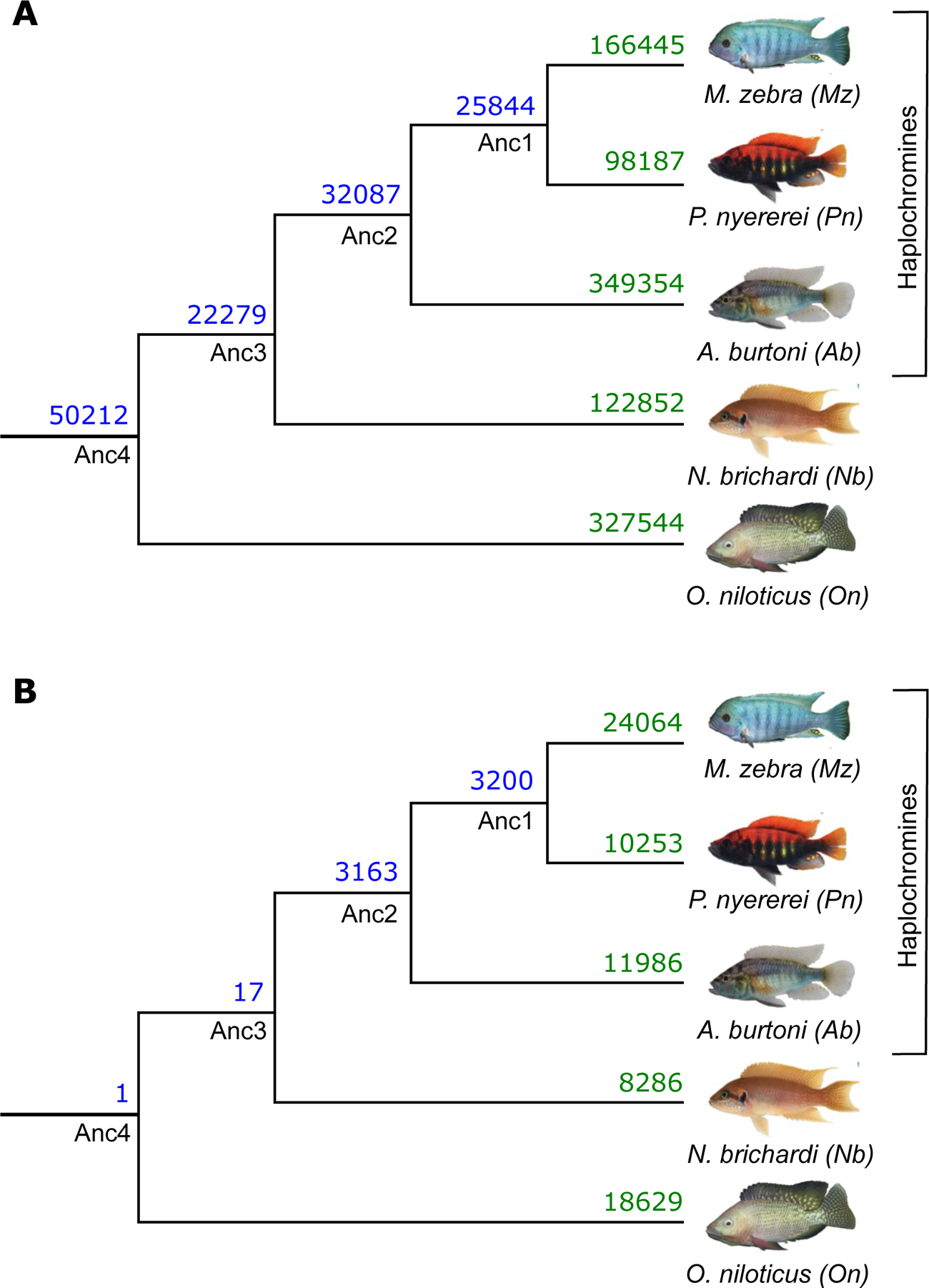
Evolution of miRNA binding sites along the five cichlid phylogeny. Number of shared and non-shared target sites based on miRNA binding site overlap in multiple 3’ UTR alignments are shown at ancestral nodes (in blue) and branches (in green) for **(A)** same miRNA families and **(B)** different miRNA families.

### Comparative analysis of three-node motifs identifies increased novel network architecture between the five cichlid species

Since a GRN can be composed of both transcriptional activation and post-transcriptional repression, we extend our previous analysis of cichlid GRN evolution (Mehta et al. 2021) by instead focusing on ‘three-node motifs’ (Alon 2007). As previously shown for mammals (Stergachis et al. 2014), the study of such motifs may serve as a reliable indicator of evolutionary conserved and diverged network signatures across species. Owing to the input dataset and our aim of focusing on the impact of miRNA associated GRN rewiring in five cichlids, we focus on a topology representative of a miRNA feed-forward loop (miRNA-FFL) (Fig. 3a). In this model, the TF is predicted to regulate a target gene (TG) and a miRNA is predicted to directly regulate either the TF or TG (Fig. 3a). According to this model and to avoid any bias of gene/miRNA loss or mis-annotations in motif/binding site comparisons across all species, we filtered a starting set of 37,320,950 three-node motif edges (Supplementary Fig. S6) for 1-to-1 orthologous TFs, TGs and miRNA families. This resulted in a final set of 17,987,294 three-node motif edges across the five species (see *Supplementary information* and Supplementary Fig. S7). In this set, 467,279 (3%) three-node motif edges are conserved across all five species (Supplementary Fig. S8, Supplementary Table S2). Instead, 1,321,875 (7%) – 3,124,263 (17%) three-node motifs are unique to each species (Supplementary Fig. S8, Supplementary Table S2). In the 17,987,294 three-node motif edges, we identified 429,197 (TF:TG) and 366,302 miRNA:TG unique edges across the five species. Using a presence and absence matrices of these unique edges, we note that on average, 56% of miRNA:TG edges are lost compared to 46% of all TF:TG edges across five species (see *Supplementary information* and Supplementary Fig. S9).

**Fig. 3.**
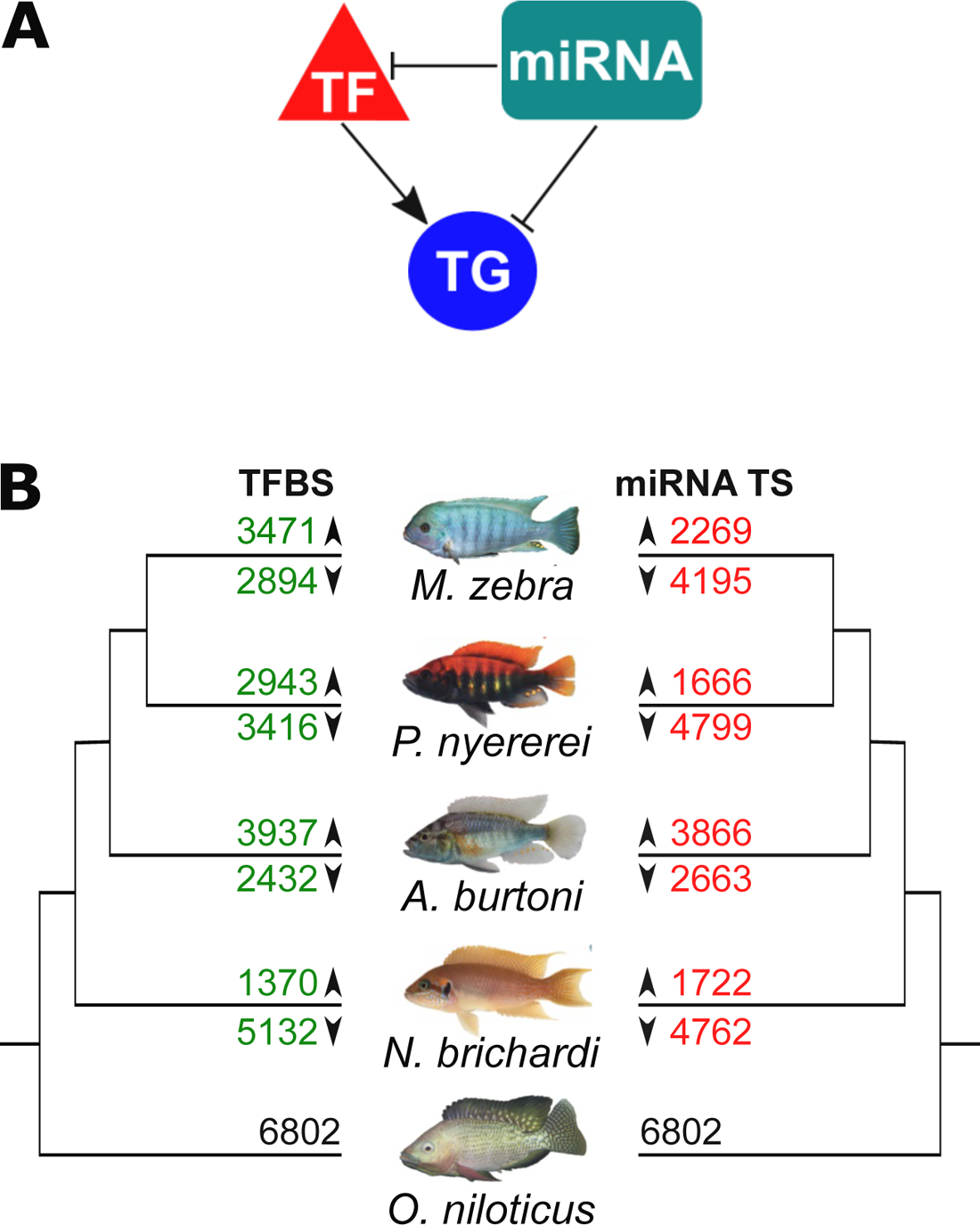
Evolution of three-node motifs (TF-TG-miRNA) in cichlids. **(A)** Three-node motif model used to assess network architecture. The three-node motif model used is representative of a miRNA feed-forward loop (miRNA-FFL). TF – Transcription Factor; TG – Target Gene. **(B)** TFBS and miRNA target site gain and loss in edges of 1-to-1 orthogolous target genes in three-node motifs of four cichlids. Five cichlid phylogeny showing number of 1-to-1 target gene orthologs with either TFBS (on left, in green) or miRNA (on right, in red) gain (above branch) or loss (below branch) vs *O. niloticus*. Binding sites in *O. niloticus* were used as reference for calculating gains and losses in the other species for 1-to-1 orthogroups (black numbers).

Using the same unique edges of each TG, we identified 115,031 unique TF:miRNA relationships and assessed their frequency to identify co-regulatory conservation and divergence along the phylogeny (see *Materials and Methods*). Of these TF:miRNA relationships, 25,209 (22%) are conserved across all five species. An example of one such conserved relationship is *miR-18*, a miRNA with negatively correlated expression with mRNA pairs in Midas cichlids (Franchini et al. 2019), being paired with NR2C2, a TF that we previously implicated in visual opsin GRN rewiring in cichlids (Mehta et al. 2021). On the other hand, 35,137 (31%) TF:miRNA relationships are unique to any one species and target an average of 5,658 genes across the five species, of which, 25 out of the 90 genes associated with phenotypic diversity from previous studies are also targeted (Supplementary Table S3, see *Supplementary information*). For example, *fgfr1,* a gene implicated in shaping cichlid scales (Albertson et al. 2018), is a species-specific target in *A. burtoni* of 48 co-regulatory relationships e.g. KLF5B:*miR-27e*; and IRF7:*miR-27b* is a unique co-regulatory relationship of *M. zebra,* and targets the fast-evolving (Brawand et al. 2014) morphogenesis gene, *bmpr1* (Supplementary Fig. S10, see *Supplementary information*). Overall, by looking at three-node motifs, we identify evolutionary conserved signatures as well as much more novel species-specific network architecture that can be associated with traits of cichlid phenotypic diversity.

### Network rewiring is associated with different models of regulatory binding site turnover in three-node motifs across species

The previous section focused on the evolution of whole network motifs. Here, we determine whether species differences in edges of these motifs are due to regulatory binding site turnover associated with previously described GRN rewiring events (Mehta et al. 2021). Using the unique TF:TG (429,197) and miRNA:TG (366,302) edges of 6,802 1-to-1 TG orthogroups, we note variation in TF or miRNA binding site gain or loss along the phylogeny (Fig. 3b). In the haplochromines, both *M. zebra* (4,195) and *P. nyererei* (4,799) have more TGs with miRNA binding site losses, whereas in *A. burtoni*, there are more TGs with either TFBS (3,937) or miRNA (3,866) gain (Fig. 3b). On the other hand, *N. brichardi* has more TGs (5,132) with TFBS and miRNA binding site loss (Fig. 3b).

We then sought to test the impact of miRNA binding site turnover in the three-node motifs and characterise the model of binding site evolution. It also provides us with the opportunity to assess the relative contributions of miRNA and TF binding sites turnover to previously observed GRN rewiring events (Mehta et al. 2021). We previously measured rewiring rates of TFBSs (Mehta et al. 2021) using DyNet (Goenawan et al. 2016), whereby the variance of nodes and TF-TG edges in orthologous gene networks is calculated, and a rewiring metric score (degree-corrected *D_n_* score) is outputted (Goenawan et al. 2016). After ordering the *D_n_* score, calculating the mean for all orthogroups, and testing the significance of difference around the mean, a degree-corrected *D_n_* score >0.17 was characterised as a threshold for significant GRN rewiring (Mehta et al. 2021). To test the associations of GRN rewiring and binding site turnover, we use and extend our analyses in Fig. 3b whereby *O. niloticus* 1-to-1 orthologous genes are used as a reference to assign each of the other species genes to one of eight models of binding site evolution, including all combinations of TFBS/miRNA gain, loss, or ‘no change’. We then tested the significance of enrichment (hypergeometric *p*-val <0.05) of orthogroups in each model of binding site evolution that could be contributing to either 6,542 significantly rewired (degree-corrected *D_n_* score >0.17) or 260 low to non-rewired (degree-corrected *D_n_* score ≤0.17) 1-to-1 orthogroups (see *Materials and Methods*). We report that TFBS gain/loss (mean degree-corrected *D_n_* score = 0.21), instead of miRNA binding site gain/loss (mean degree-corrected *D_n_* score = 0.20), had the largest effect on significantly rewired (degree-corrected *D_n_* score >0.17) orthologs (Fig. 4a, Supplementary Fig. S11). The most associated models of rewired orthologs are TFBS loss in *A. burtoni* (*p*-value =0.009) and TFBS gain in *M. zebra, P. nyererei* and *N. brichardi* (*p*-value =0.0007-0.03) (Supplementary Table S5). However, all low to non-rewired orthologs (*D_n_* score ≤ 0.17) that should be impervious to TFBS-based rewiring, are expectedly most associated with no change in TFBS, but miRNA binding site loss in all four species (*p*-value =0.000005-0.05) (Supplementary Table S6). This therefore indicates a discrete impact of GRN rewiring based on miRNA binding site loss. Overall, this suggests that different models of regulatory binding site evolution have impacted GRN rewiring in the studied cichlid lineages.

**Fig. 4.**
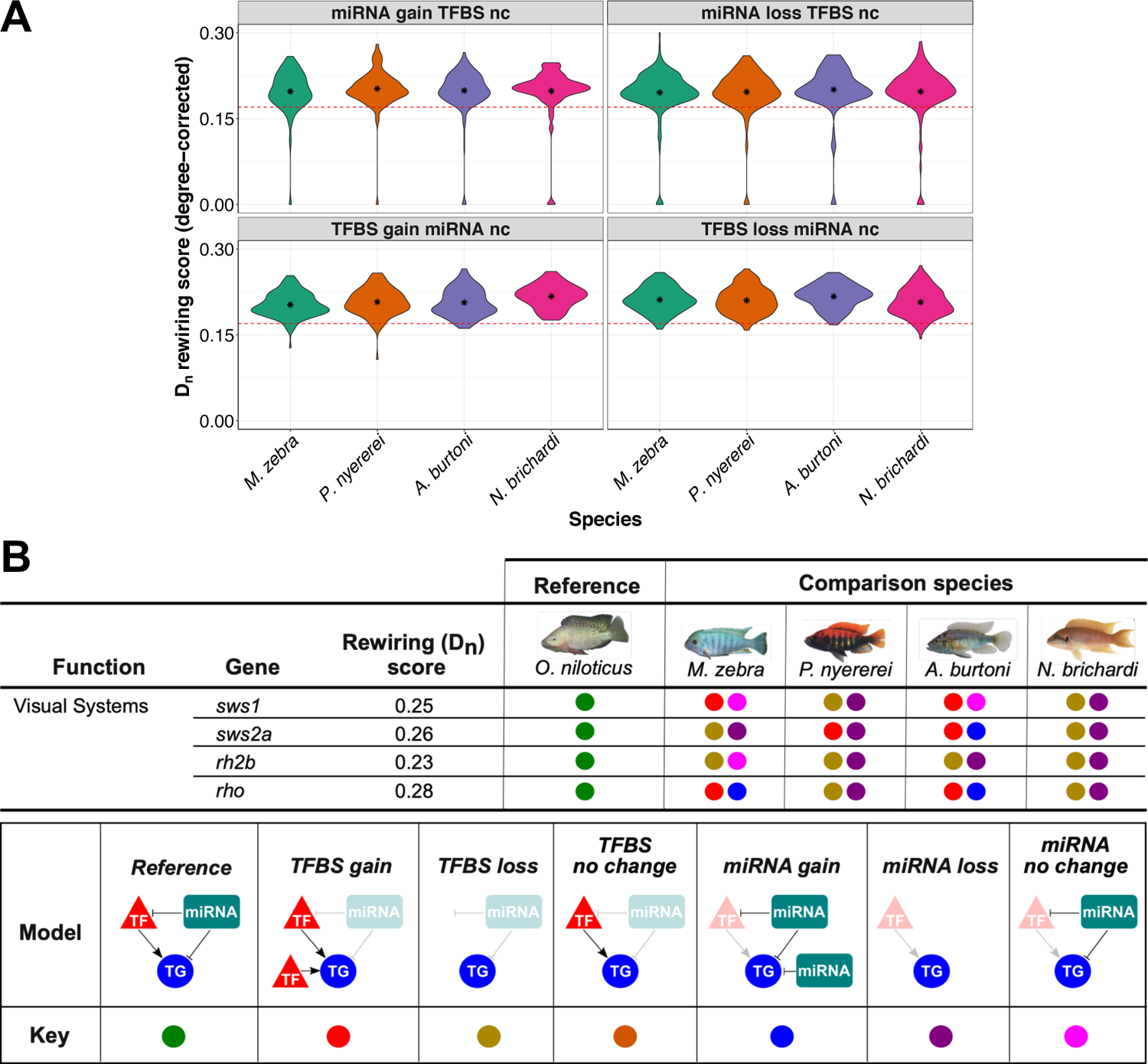
Binding site evolution in three node motifs of cichlid genes and their association with rewiring events. **(A)** Different models of TFBS and miRNA binding site evolution with associated rewiring rates of 1-to-1 orthogroups in four cichlids. Violin plots of 4/8 models of binding site evolution in each species (x-axis) with DyNet rewiring score of each 1-to-1 orthogroup as degree corrected D_n_ score (y-axis). Red dotted line demarcates a *D_n_* score threshold of 0.17 (for rewired vs low to non-rewired genes), which was set based on the mean *D_n_* score for all orthogroups in our previous study (Mehta et al. 2021). The term ‘nc’ refers to no change and mean values are shown as internal asterisk. All statistics are included in Table S5 and violin plots of other models in Fig. S11. **(B)** Binding site evolution of four cichlid visual system genes. DyNet rewiring (*D_n_*) score for all genes obtained from our previous study (Mehta et al. 2021). For the four comparison species, each genes model of TFBS and miRNA target site evolution in three-node motifs is calculated using the orthologous *O. niloticus* gene as a reference and demarcated as per the ‘model’ and ‘key’ in legend. All statistics are included in Supplementary Table S4-5.

### Regulatory binding site turnover in three-node motifs is associated with network rewiring of adaptive trait genes

Further examination of the rewired orthologs with either of the eight models of binding site evolution identifies teleost and cichlid trait genes associated with phenotypic diversity from previous studies (Fig. 4b, Supplementary Fig. S13, Supplementary Table S3, see *Supplementary information*). Compared to all orthologs (mean *D_n_* rewiring score = 0.17), we previously showed that four visual opsin genes (*sws1*, *rho*, *sws2a*, and *rh2b*) have considerably rewired networks (*D_n_* score = 0.23-0.28) in species utilising the same wavelength visual palette and opsin genes (Mehta et al. 2021). The evolution of GRNs and utilisation of diverse palettes of co-expressed opsins is able to induce shifts in adaptive spectral sensitivity of adult cichlids (Carleton 2009). Such quantitative and qualitative changes of opsin gene expression can fine-tune cichlid vision in response to prey, ambient light changes, or even conspecific signals and is thought to be primarily achieved through differential regulation (Hofmann et al. 2009; O’Quin et al. 2012). In this respect, we previously demonstrated that opsin expression diversity could be the result of TF regulatory divergence in cichlids (Mehta et al. 2021). By investigating binding site evolution in our three-node motifs, we are able to further identify the genetic factors that could be associated with the regulation of opsin expression variation between species. The *sws1* (ultraviolet-sensitive) opsin, utilised as part of the short-wavelength palette in *M. zebra* and *N. brichardi*, has TFBS gain and no change in miRNA binding site in *M. zebra* and *A. burtoni*, but TFBS and miRNA binding site loss in the other two species (Fig. 4b). In another example, *rhodopsin* (*rho*), associated with dim light vision in all species, has TFBS and miRNA binding site gain in *M. zebra* and *A. burtoni*, but TFBS and miRNA binding site loss in the other two species (Fig. 4b). These patterns of TF and miRNA regulatory divergence, including that of other visual opsins e.g. *sws2a* and *rh2b* (Fig. 4b), could therefore contribute to differential expression of adaptive trait genes (see *Supplementary information*), including visual opsins and their fine-tuning.

### Discrete changes at regulatory sites are fast-evolving and associated with binding site turnover

To study the evolution of TF and miRNA regulatory divergence in the five cichlids, we assessed whether regulatory binding site turnover in three-node motifs is occurring at regions with a different rate of evolution than that expected under a neutral model. We did this by 1) determining the rate of evolution at fourfold degenerate sites and regulatory regions (3’ UTR, up to 5kb gene promoter, miRNA binding sites and TFBSs); 2) identifying between species variation at regulatory sites and test for accelerated evolution; and 3) assessing corresponding regions in the context of phylogeny and ecology of radiating lake species. We started with 20,106 - 24,559 (3’ UTR), 19,706 – 24,123 (up to 5kb gene promoter), 232,050 – 478,796 (miRNA binding sites), and 3,790,407 – 7,064,048 (TFBSs) unique regulatory regions across the five species, and as a putatively neutrally evolving comparison, 5,292,087 – 6,539,362 fourfold degenerate sites (Supplementary Table S9). The rate of substitutions in whole genome pairwise comparisons was calculated using phyloP (Pollard et al. 2010). In total, 86 - 98% of the nucleotides investigated had mapped conservation-acceleration (CONACC) scores (Supplementary Table S9). Across all five species pairwise comparisons, 92% of the fourfold degenerate sites are conserved, which is consistent with an average of ∼6% pairwise divergence at fourfold sites between *O. niloticus* and the other four species (Brawand et al. 2014), whereas 3% are evolving at a faster rate than that expected (Supplementary Fig. S14, Supplementary Table S10). On the other hand, 81% of the regulatory regions are conserved, and 4% are exhibiting accelerated evolution (Supplementary Fig. S14, Supplementary Table S10). Since our previous study found that discrete regulatory mutations are driving GRN rewiring events (Mehta et al. 2021), we hypothesised that such mutations could account for some of the accelerated regulatory sites. Using pairwise polymorphic nucleotide sites in each of the four regulatory regions (Supplementary Table S12), we identified that 81-87% (3’ UTR), 69-77% (up to 5kb gene promoter), 83-99% (miRNA binding sites), and 6-8% (TFBSs) of accelerated sites are accounted for by variation in a single species (Supplementary Fig. S15, Supplementary Table S14). Notably, the proportion of these accelerated sites in the regulatory regions are significantly different (Wilcoxon rank sum test, adjusted *p-*value <0.05, see *Materials and Methods*), especially between TF and miRNA binding sites both within, and between species (Supplementary Table S15). These results support the notion that discrete mutations in TFBSs (Mehta et al. 2021), albeit it very few, and in miRNA binding sites are fast evolving i.e. fast-evolving regulatory mutations, and drive regulatory binding site turnover in three-node motifs of the five cichlids.

### Discrete changes at regulatory sites are associated with regulatory binding site turnover in adaptive trait genes

Our previous study identified an abundance of adaptive trait genes with comparatively higher rewired (*D_n_* score >0.17) networks (based on TFBSs), compared to all orthologs (Mehta et al. 2021). As a measure of regulatory binding site turnover, we therefore sought to test the frequency of association of fast-evolving regulatory mutations in 90 adaptive trait genes (Supplementary Table S3) compared to those in corresponding regulatory regions of 90 random ‘no to low rewired’ genes (*D_n_* score ≤ 0.17) from our previous study (Mehta et al. 2021) (see *Supplementary information*). We used the ‘no to low rewired’ genes to ensure that the test is not biased towards genes that have rewired GRNs based on TF divergence and using the Wilcoxon rank sum test, tested the frequency 1000 times to ensure sufficient randomisation of ‘no to low rewired’ (see *Materials and Methods*). By comparing the proportion of fast-evolving regulatory mutations in corresponding regions of 90 adaptive trait genes and 90 random no to low rewired genes, the most notable differences (>950/1000 Wilcoxon rank sum tests, adjusted *p-value* <0.05) are found in the proportion of accelerated nucleotides in TFBSs of 90 adaptive trait gene promoter regions (Supplementary Table S16). We identified 17 adaptive trait genes with significant turnover between TF and miRNA binding sites (Supplementary Table S17-18). In *M. zebra, P. nyererei,* and *O. niloticus*, this includes genes associated with brain development and neurogenesis e.g. *neurod1*, morphogenesis e.g. *bmpr1*, and visual opsins e.g. *rho* and *sws1* (Supplementary Table S17).

Furthermore, fast evolving regulatory mutations of miRNAs and TFs could be associated with the function of adaptive trait genes like, for example, ATF3 associated with neuroprotection of the retina (Kole et al. 2020) and *miR-99* implicated in retinal regulatory networks (Andreeva and Cooper 2014) are both predicted to target the visual opsin *sws1*, and MXI1 associated with neurogenesis (Klisch et al. 2006) and *miR-212* associated with synaptic plasticity and function (Remenyi et al. 2013) is predicted to target the dim-light visual opsin, *rho* (Supplementary Table S18). Discrete mutations in regulatory binding sites of cichlid adaptive trait genes could therefore be driving GRN evolution associated with traits of cichlid phenotypic diversity.

### Discrete changes at regulatory regions of adaptive trait genes segregate according to phylogeny and ecology of radiating cichlids

In our previous study, we identified that discrete TFBS mutations driving GRN evolution of visual opsin genes, also segregate according to the phylogeny and ecology of radiating lake species (Mehta et al. 2021). Here, we extend this approach to study both TF and miRNA binding site variation of three-node motifs in the context of phylogeny and ecology of lake species. Using the Lake Malawi species, *M. zebra,* as a reference, we assess whether regulatory binding site turnover in three-node motifs of this species could be genotypically associated with the ecology of sequenced Lake Malawi species (Malinsky et al. 2018). For this, we started with 827 nucleotide sites that 1) have identified variation between *M. zebra* and any of the other four cichlid species; 2) are located in binding sites of either TFs (709 nucleotide sites) or miRNAs (118 nucleotide sites) of *M. zebra* adaptive trait genes, that also have a significant difference (adjusted *p-value* <0.05) in the proportion of accelerated nucleotides, indicative of regulatory binding site turnover in their associated three-node motifs; and 3) are evolving at a significantly faster rate (adjusted *p-value* <0.05) than expected under a neutral model (Supplementary Table S18). We identified that 94 out of 827 accelerated nucleotide sites with between species variation across 73 Lake Malawi species, also exhibit flanking sequence conservation, representative of shared ancestral variation. Of the 94 accelerated nucleotide sites, 21 are found in miRNA binding sites, and 73 are found in TFBSs of which, 55 were not identified in our previous study (Mehta et al. 2021) due to not incorporating substitution rates.

Amongst the 76 accelerated nucleotide sites uniquely identified in this study, 15 (20%) include TF and miRNA binding site variation of visual opsin genes. Given the variability and importance of visual systems towards cichlid foraging habits, we therefore focus on variation at accelerated regulatory regions of visual opsin genes. If the TF and miRNA binding sites are likely functional, we hypothesise that radiating species with similar foraging habits would share conserved regulatory genotypes, to possibly regulate and tune similar spectral sensitivities; whereas distally related species with dissimilar foraging habits would segregate at the corresponding regulatory site.

We first focus on a three-node motif of the *M. zebra* short wavelength palette visual opsin gene, *sws1*, that is predicted to be regulated by *miR-99a* and ATF3 (Fig. 5a). The homozygous variant (C|C) that predicts binding of *miR-99a* to *M. zebra sws1* 3’ UTR (Fig. 5a) is 1) conserved in 60/134 (45%) Lake Malawi individuals, including the fellow algae eater, *T. tropheops,* and other distantly related species e.g. *D. kiwinge* and *N. polystigma*, that utilise the same short wavelength palette; but 2) lost in the other four species due to the A/A homozygous variant (Fig. 5a) and also homozygous segregated (A|A) in 38/134 (28%) Lake Malawi individuals, including its most closely related Mbuna species *(P. genalutea*) and *A. calliptera* (Fig. 5b and Supplementary Fig. S17). Another homozygous variant (C|C), that predicts binding of ATF3 to *M. zebra sws1* gene promoter, but is lost in *O. niloticus,* due to the T/T homozygous variant (Fig. 5a), is 1) conserved in all closely related Mbuna species and 102/116 (88%) Lake Malawi individuals, including the closely related *A. calliptera* clade; but 2) heterozygous or homozygous segregating in distantly related Lake Malawi species that utilise the same short wavelength palette, but occupy different habitats and foraging habits e.g. *D. kiwinge* – T|T and *N. polystigma* – T|C (Fig. 5b and Supplementary Fig. S16). Overall, this suggests that whilst *miR-99a* could be core regulator of *sws1* in nearly half of the studied Lake Malawi species, it is 1) unlikely to be a co-regulator of *sws1* (with ATF3) in either distantly related Lake Malawi species utilising the short wavelength palette e.g. *D. kiwinge* and *N. polystigma*, or the *A. calliptera* clade; but 2) likely to co-regulate *sws1* (with ATF3) in most members of the rock-dwelling Mbuna clade (Fig. 5b and Supplementary Fig. S16-17). In another example, we show that a three-node motif of the dim-light vision gene, *rho,* consisting of *miR-212* and MXI1 has conserved regulatory genotypes in all studied Lake Malawi species, but has segregated and therefore not predicted in the other four cichlids (Supplementary Fig. S18-19). Phylogenetic independent contrast analysis (Felsenstein 1985) of the *sws1* (Supplementary Fig. S20-21) and rho (Supplementary Fig. S22-23) genotypes against visual traits and ecology of each of the 73 Lake Malawi species, highlights very little change in correlation once the phylogeny is taken into account and a regression model fitted (see *Materials and Methods*). In summary, we identified three-node motifs of visual systems that segregate according to phylogeny and ecology of lake species. Regulatory binding site turnover of three-node motifs is therefore a key contributing mechanism of GRN evolution associated with adaptive innovations in East African cichlid radiations.

**Fig. 5.**
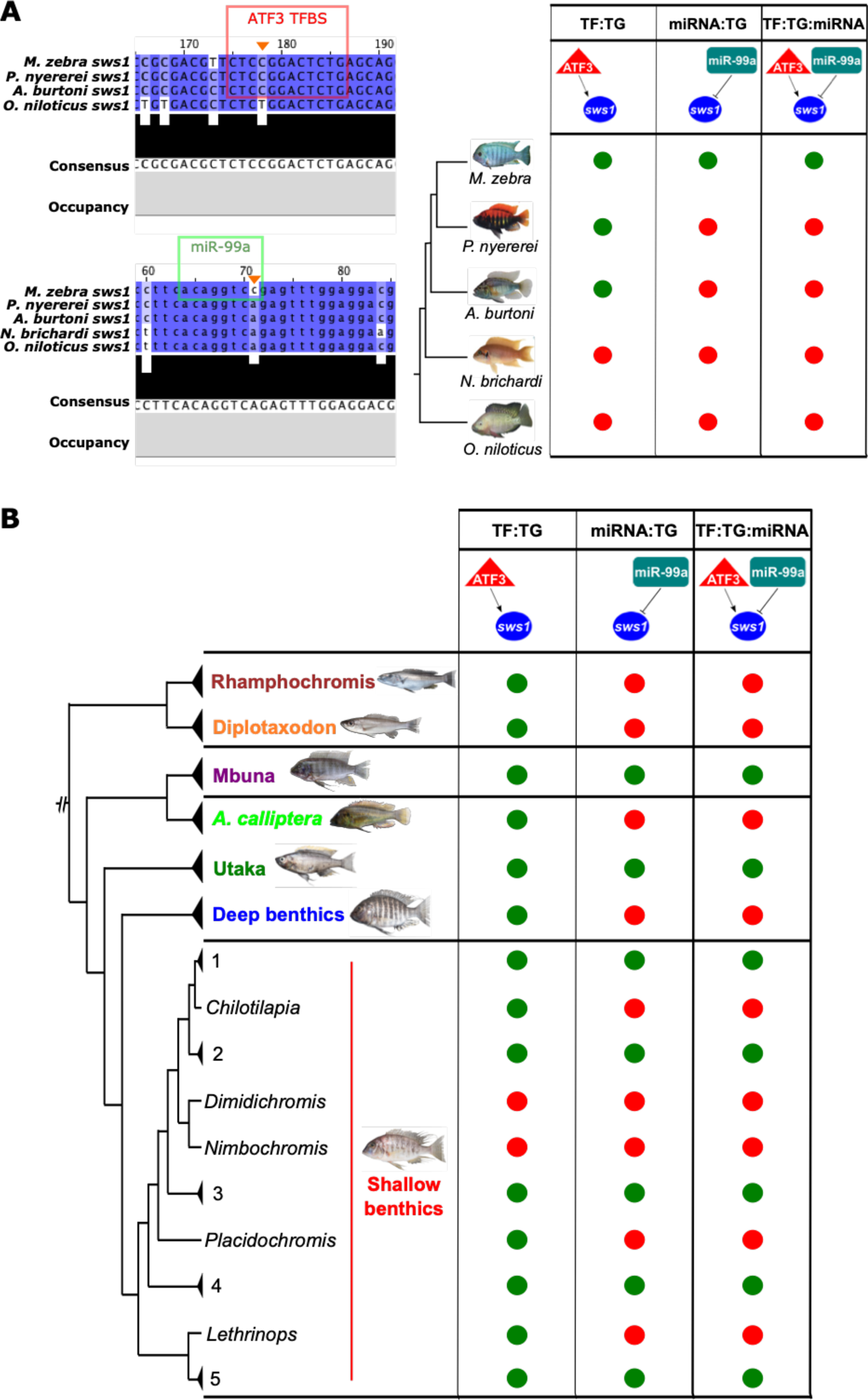
Evolution of the ATF3:*sws1*:miR-99a three-node motif in the five cichlids and Lake Malawi species. **(A)** On the *top left,* ATF3 motif prediction in *M. zebra, P. nyererei* and *A. burtoni sws1* gene promoter (red box) and substitution demarcated in *O. niloticus sws1* gene promoter (orange arrow). On the *bottom left,* miR-99a binding site prediction in *M. zebra sws1* 3’ UTR (green box) and substitution demarcated in *P. nyererei, A. burtoni, N. brichardi* and *O. niloticus sws1* 3’ UTR (orange arrow). According to these predicted sites, evolution of the ATF3:*sws1*:miR-99a three-node motif in the five cichlid phylogeny is depicted based on presence (green circle) and absence (red circle). **(B)** Simplified presence (green circle) and absence (red circle) of the ATF3:*sws1*:miR-99a three-node motif in Lake Malawi species based on SNP genotypes overlapping ATF3 TFBS and miR-99a binding sites in *M. zebra sws1* gene promoter and 3’ UTR (orange arrows, Fig.5a). Lake Malawi phylogeny reproduced from published ASTRAL phylogeny (Malinsky et al. 2018). Phylogenetic branches labelled with genus, species or clade identifiers. Within the shallow benthics, species within some clades are summarised by numbers: 1 – *Hemitaeniochromis, Protomelas*; 2 – *Hemitilapia, Otopharnyx, Mylochromis*; 3 – *Champsochromis, Tyrannochromis, Trematocranus, Otopharnyx, Mylochromis, Stigmatochromis, Taeniochromis, Buccochromis, Ctenopharynx;* 4 – *Mylochromis;* 5 – *Taeniolethrinops.* Expanded genotype, phenotype and ecotype phylogeny in Supplementary Fig. S16-17.

## Discussion

Evolutionary changes of regulatory systems and GRN rewiring events can contribute to the evolution of phenotypic diversity and rapid adaptation (Kratochwil and Meyer 2015). This is particularly the case for East African cichlid diversification that has been shaped by complex evolutionary and genomic forces. These include divergent selection acting upon regulatory regions that can alter gene expression programmes (Brawand et al. 2014), rapid evolution of noncoding RNA expression (El Taher et al. 2021) and ancient polymorphisms in noncoding regions (McGee et al. 2020), contrasted against a background of low between-species genetic diversity (Malinsky et al. 2018; McGee et al. 2020; Ronco et al. 2021). All of these findings imply that discrete differences at regulatory regions could contribute to phenotypic differences and indeed, through discrete changes in TFBSs, we previously showed that GRN rewiring could be a key contributor to cichlid phenotypic diversity (Mehta et al. 2021). However, our previous study did not explore and integrate other genetic mechanisms, like the contributions of miRNAs towards cichlid GRN evolution. Given that miRNAs are key regulators of post-transcriptional gene expression, and that novel miRNAs have evolved in rapidly radiating cichlids (Brawand et al. 2014; Franchini et al. 2016; Franchini et al. 2019; Xiong et al. 2019), they could therefore contribute to GRN evolution associated with cichlid phenotypic diversity.

Across the five cichlid species, we identified a total of 15,390,993 binding sites for the 18,799 co-expressed orthogroups. The total number of sites is inflated compared to the 38,768 putative miRNA binding sites predicted in the Midas genome (Franchini et al. 2016) due to 1) a difference of approaches whereby, Franchini *et al*. use miRanda (John et al. 2004) for target prediction, a tool that is found to predict fewer experimentally derived sites as compared to Targetscan7 (Agarwal et al. 2015; Riffo-Campos et al. 2016); 2) retainment of multiple sites for the same miRNA along a 3’ UTR to analyse nucleotide variation at all predicted sites, like that previously applied for all TFBSs in corresponding gene promoter regions (Mehta et al. 2021); and 3) we predict binding sites in an average of 13,998 3’ UTRs, which is 41% more (8,232 3’ UTRs) than in the Midas genome (Franchini et al. 2016). Despite these differences, we identify an average of 356,379 unique miRNA binding sites across the five species, ranging from 1 to 173 unique miRNA binding sites per 3’ UTR, that is comparable to the range (0-222) previously identified in Midas cichlids (Franchini et al. 2016). Across the five species, 3’ UTR binding sites are differentially predicted for up to 172 miRNA families. The under-representation of certain families in a species can be attributed to mutations of the seed sequence and *arm* switching (Berezikov 2011; Brawand et al. 2014). The largest number of conserved miRNA families are across all five species and include binding sites in 3’ UTRs of genes associated with jaw development (Bloomquist et al. 2017) and deep-water adaptation (Hahn et al. 2017). This supports an important regulatory role of miRNAs to cichlid adaptive traits (Brawand et al. 2014) over a divergence time of ∼ 19 MYRs (Hughes et al. 2018). We identified more miRNA family divergence within the haplochromine lineage, particularly in 3’ UTRs of developmental genes; a finding that is consistent with rapid evolutionary changes of noncoding RNA expression (El Taher et al. 2021) and noncoding regions (McGee et al. 2020) in corresponding Lake Tanganyika and Victoria species. Our results suggest a deeply conserved role of miRNA regulation in the five cichlids however, binding site divergence of miRNA families is likely to have an important gene regulatory role in the rapid (∼ 6 MYRs (Hughes et al. 2018)) phenotypic divergence of haplochromines.

Since a GRN can be composed of both transcriptional activation and repression, we extended our previous study (Mehta et al. 2021) to focus on a miRNA feed-forward loop ‘three-node motif’. Using this three-node motif as a measure of network divergence and evolutionary constraint, we identified increased novel/species-specific three-node motifs overall, reflected by a higher rate of miRNA edge loss (than TF edge loss) along the phylogeny. This is consistent with previous findings in Midas cichlids where miRNAs and concomitantly, their binding sites, can be rapidly lost between related groups (Xiong et al. 2019). In support, we tested the association of eight models of TFBS and/or miRNA binding site evolution, including ‘no change’, on TG edges previously defined as low to non-rewired (Mehta et al. 2021) based on TFBSs only. We found that the most associations were expectedly with no change in TFBS, and miRNA binding site loss in all four species compared to *O. niloticus* as a reference. This indicates that miRNA binding site loss is having a discrete impact on GRN rewiring, but overall, different models of regulatory binding site evolution have impacted GRN rewiring in the cichlid lineages studied here. This included identifying that the most associated model of four highly rewired visual opsin genes (*sws1*, *rho*, *sws2a*, and *rh2b*) (Mehta et al. 2021) was generally TFBS (in 50%) and miRNA binding site (in 66%) loss across the species. This supports our previous work demonstrating that opsin expression diversity could be the result of TFBS divergence in cichlids (Mehta et al. 2021) and thus, regulatory divergence is likely to accommodate for heterochronic shifts in opsin expression (Carleton et al. 2008; O’Quin et al. 2011). Overall, these findings suggest that differential patterns of TF and miRNA regulatory divergence are likely to be associated with three-node motif and GRN rewiring of cichlid adaptive traits.

Across all five species pairwise comparisons, we find that regulatory divergence i.e. binding site turnover in three-node motifs is occurring at regions with a different rate of evolution than that expected under a neutral model. This is supported by a previous study that also identified evolutionary-accelerated 3’ UTRs in the same five cichlid species and overall, suggested this as a contributory mechanism for speciation (Puntambekar et al. 2020). However, we extend all previous work to show that on average, nearly a third of all fast-evolving nucleotide sites in the four regulatory regions (3’ UTR, up to 5kb gene promoter, miRNA binding sites and TFBSs) can be explained by pairwise polymorphisms in a single species. Whilst more than 83% of fast-evolving nucleotides in miRNA binding sites are accounted for variation in a single species, less than 8% of TFBSs are accounted for by the same type of fast-evolving variation. This supports our previous finding of discrete mutations in TFBSs driving GRN rewiring events (Mehta et al. 2021), as well as elevated SNP densities in predicted miRNA binding sites, compared to flanking 3’ UTR regions, of five Lake Malawi species (Loh et al. 2011). Positive selection acting upon these regulatory regions is therefore likely to be an important evolutionary force in rapidly radiating cichlids. This is especially the case for adaptive trait genes such as the visual opsins e.g. *rho* and *sws1*, that we show to exhibit a higher proportion of fast-evolving nucleotides in their TF and miRNA binding sites, compared to subsets of random genes. Furthermore, these TFs and miRNAs are generally functionally associated with their target gene in predicted three-node motifs like, for example, the visual opsin gene, *sws1*, is predicted to be co-regulated by the TF, ATF3, that is associated with neuroprotection of the retina (Kole et al. 2020) and *miR-99* implicated in retinal regulatory networks (Andreeva and Cooper 2014). The regulatory variants of this three-node motif (ATF3 > *sws1* < *miR-99a*) in *M. zebra* also appear to differentially segregate according to phylogeny and ecology of Lake Malawi species (Malinsky et al. 2018). We find that ATF3:*miR-99a* could be an important regulator of *sws1* in the rock-dwelling Mbuna clade, but unlikely to co-regulate *sws1* as part of the short-wavelength palette in the *A. calliptera* clade and distantly related Lake Malawi species. For another opsin gene, we identified that the possible neural co-regulation of *rho*, and therefore dim-light vision response by MXI1:*miR-212*, could be a Lake Malawi specific regulatory innovation. Overall, differential binding of miRNAs and TFs associated with retinal sensory modalities (Loh et al. 2011) and visual tuning (Sandkam et al. 2020) is likely to be an important genetic mechanism contributing to Lake Malawi species visual adaptations. Whilst these results significantly expand our previously characterised visual opsin GRNs (Mehta et al. 2021) and provide insights into their evolution in radiating cichlids, we also provide support for the hypothesis that the evolution of cichlid visual tuning has been facilitated by regulatory mutations that are constrained by mutational dynamics (Nandamuri et al. 2018; Sandkam et al. 2020). Differential regulation of opsin genes in three-node motifs between cichlid species and their implications towards visual tuning could correspond to diversity of foraging habits, diet, habitat choice and also nuptial colouration. Fitting the Lake Malawi phylogeny had little effect on the correlations between regulatory genotypes, and visual/ecological characteristics, and therefore suggests covariance between TF/miRNA regulatory genotypes and traits. However, similar to our previous study (Mehta et al. 2021), weak correlation suggests that ecotype-associated three-node motif and GRN rewiring requires additional testing. This analysis would further benefit from 1) supplementing any missing data (of wavelength palette, habitat and/or foraging habit/diet); 2) adding species data from any lowly represented clades e.g. Mbuna; and 3) experimental testing of the predicted sites.

Alongside our previous study (Mehta et al. 2021), the three-node motifs and extended GRNs generated here represent a unique resource for the community; powering further molecular and evolutionary analysis of cichlid adaptive traits. For example, further examination of the three-node motifs predicted for the visual systems, that could co-regulate opsin expression diversity, could further shed light on previous preliminary studies (Carleton et al. 2008; Hofmann et al. 2009; O’Quin et al. 2012; Nandamuri et al. 2018; Sandkam et al. 2020). This could involve functional validations of three-node motifs to observed trait variation by 1) high-throughput miRNA-mRNA complex and protein-DNA assays to confirm binding of thousands of sites; 2) reporter and/or cell-based assays to demonstrate transcriptional regulation; and 3) genome editing e.g. CRISPR mutations of regulatory variants to test for any observed phenotypic effect. Nonetheless, by studying the impact of miRNA regulation in three-node motifs, this work extends the first genome-wide exploration of GRN evolution in cichlids (Mehta et al. 2021). In a wider context, as the individual regulatory hallmarks of TFs and miRNAs start to become characterised in disease e.g. forms of cancer (Plaisier et al. 2016; Mullany et al. 2018; Nersisyan et al. 2021), congenital heart disease (You et al. 2020), neuromuscular disorders (Bo et al. 2021), as well as related to gene expression in human tissues (Minchington et al. 2020) and plant stress response (Sharma et al. 2020; Sharma et al. 2021), the computational framework we applied here could be used to study the evolution of characterised regulatory edges and GRNs in the aforementioned, and other systems and phylogenies. However, the combined framework could be extended further by 1) analysing the impact of either more, or all of the 104 three-node motif models (Ahnert and Fink 2016) through the integration of epigenetic and co-immunoprecipitation assay data to gain regulatory directionality; and 2) including relevant datasets to study the regulatory effect of other mechanisms e.g. lncRNAs and enhancers on network topology, that could also contribute towards the evolution of cichlid phenotypic diversity (Brawand et al. 2014; Salzburger 2018). Whilst many of the predicted three-node motifs could be false positives, the approach applied here and previously (Mehta et al. 2021) ensured for rigorous filtering at each step; this included stringent statistical significance measures, and all whilst accounting for any node loss and mis-annotations in selected species (see *Materials and Methods*).

In summary, cichlids appear to utilise an array of genetic mechanisms that also contribute towards phenotypic diversity in other organisms (Wittkopp et al. 2008; Yanai and Hunter 2009; Chan et al. 2010; Jones et al. 2012; Thompson et al. 2013; Ichihashi et al. 2014). However, here we provide support of TF and miRNA co-regulatory rewiring in three-node motifs of genes associated with adaptive traits in cichlids. This is further supported by large-scale genotyping studies of the predicted regulatory sites in rapidly radiating cichlid species (Malinsky et al. 2018). This potential link between the evolution of three-node motifs as part of GRNs associated with cichlid adaptive traits requires further experimental verification. This is beyond that described for *cis*-regulatory sites previously (Mehta et al. 2021), as well as support based on large-scale genotyping (Malinsky et al. 2018; McGee et al. 2020; Ronco et al. 2021) and transcriptome evolution (El Taher et al. 2021); epigenetic divergence (Kratochwil and Meyer 2014; Vernaz et al. 2021); transgenesis assays (Brawand et al. 2014; Santos et al. 2014); population studies and CRISPR mutant assays (Kratochwil et al. 2018); and transcriptomic/*cis-*regulatory assays (Hofmann et al. 2009; O’Quin et al. 2010; Nandamuri et al. 2018; Sandkam et al. 2020) of cichlid species.

## Materials and Methods

### Genomic and transcriptomic resources

Genomes and transcriptomes of the five cichlid species were obtained from NCBI and corresponding publication (Brawand et al. 2014): *P. nyererei -* PunNye1.0, NCBI BioProject: PRJNA60367; BROADPN2 annotation; *M. zebra -* MetZeb1.1, NCBI BioProject: PRJNA60369; BROADMZ2 annotation; *A. burtoni -* AstBur1.0, NCBI BioProject: PRJNA60363; BROADAB2 annotation; *N. brichardi -* NeoBri1.0, NCBI BioProject: PRJNA60365; BROADNB2 annotation; *O. niloticus* - Orenil1.1 (NCBI BioProject: PRJNA59571; BROADON2 annotation.

### MicroRNA (miRNA) target prediction

We used the microRNAs that were previously sequenced from whole embryo for five cichlid species (*O. niloticus, N. brichardi*, *A. burtoni*, *P. nyererei* and *M. zebra*) (Brawand et al. 2014). The miRNA mature sequences and hairpin structures have been characterised as described previously (Brawand et al. 2014) and deposited in miRbase (Kozomara and Griffiths-Jones 2014). A total of 992 (On-198, Nb-183, Ab-243, Mz-185, Pn-183) cichlid miRNA mature sequences and annotated 3’ UTRs of 21,871 orthogroups (On-22411, Nb-20195, Ab-22662, Mz-21918, Pn-21599) in all five species (Brawand et al. 2014) were used for target prediction. We used TargetScan7 (Agarwal et al. 2015) to predict species-specific genes targeted by the sequenced microRNAs (miRNAs). We used *mafft-7.271* (Katoh and Standley 2013) to generate gene specific multiple alignments of the annotated 3’ UTRs across all five cichlid species. Target predictions were obtained by running TargetScan7 (v7.2) (Agarwal et al. 2015) according to the developer’s protocols with default parameters. Firstly, miRNA binding sites were predicted using the reformatted alignments and the sequenced mature miRNA sequences as input for the ‘targetscan_70.pl’ script with default parameters. Using the median branch length from each 3’ UTR alignment derived from the ‘targetscan_70_BL_bins.pl’ script and predicted miRNA binding sites, we then calculated the conserved branch length (BL) and probability of preferentially conserved targeting (PCT) for all predicted miRNA targets using the ‘targetscan_70_BL_PCT.pl’ script with default PCT and parameters as derived from https://www.targetscan.org/vert_72/vert_72_data_download/targetscan_70_BL_PCT.zip and (Agarwal et al. 2015). All seed sites were found and counted in 3’ UTR sequences using the ‘targetscan_count_8mers.pl’ script. Finally, to predict high-confidence miRNA targets, that could be as predictive as the most informative *in vivo* approaches such as crosslinking-immunoprecipitation (CLIP) (Agarwal et al. 2015), we used the miRNA mature sequences, 3’ UTR alignments, calculated BL and PCT scores and seed site locations with counts as input to the ‘targetscan_70_context_scores_cichlids.pl’ script to calculate Targetscan7 weighted context++ scores. The developers of Targetscan7 found that the context++ model was more predictive than any published model for miRNA targeting efficacy, and as predictive as the most informative *in vivo* crosslinking approaches e.g. crosslinking-immunoprecipitation (CLIP) (Agarwal et al. 2015). The weighted context++ score ranges from 1 to −1 and thus, scores with a lower negative value indicate a greater prediction of repression (Agarwal et al. 2015). Based on previous predictions of miRNA targets using Targetscan7 in vertebrates (Warnefors et al. 2017; Hu et al. 2019; Sayed and Park 2020), we selected all targets using a stringent weighted context++ score lower than −0.1 to filter out low quality predictions; these were the binding sites used for analyses. The multiple alignments of annotated 3’ UTRs and positions of predicted sites in each species were used to identify overlapping miRNA binding sites of miRNA families between species.

### Gene ontology (GO) enrichment

To assess enrichment of Gene Ontology (GO) terms in a given gene set, we use the Benjamini-Hochberg (Benjamini and Hochberg 1995) False Discovery Rate (FDR) corrected hypergeometric *P-*value (*q-*value). The background (control set) for the enrichment analysis is composed of all co-expressed genes (18,799 orthogroups) from our previous study (Mehta et al. 2021). GO terms for the five cichlids were extracted from those published previously (Brawand et al. 2014).

### Transcription factor (TF) motif scanning

To study transcription factor (TF) – target gene (TG) associations in three-node motifs, we used predicted transcription factor binding sites (TFBSs) from our previous study (Mehta et al. 2021). Briefly, we used the aforementioned published assemblies and associated gene annotations (Brawand et al. 2014) for each species to extract gene promoter regions, defined as up to 5 kb upstream of the transcription start site (TSS) of each gene. We used a combination of 1) JASPAR vertebrate motifs; 2) extrapolated cichlid-species specific (CS) Position Specific Scoring Matrices (PSSMs) (Mehta et al. 2021); and 3) aggregated generic cichlid-wide (CW) PSSMs (Mehta et al. 2021) to identify TF motifs. Using FIMO (Grant et al. 2011), the gene promoter regions of each species were scanned for each TF motif using either 1) an optimal calculated *p-value* for each TF PSSM, as calculated using the *matrix quality* script from the RSAT tool suite (Medina-Rivera et al. 2015); or 2) FIMO (Grant et al. 2011) default *p-value* (1e-4) for JASPAR (Khan et al. 2017) PSSMs and PSSMs for which an optimal *p-value* could not be determined. Statistically significant TFBS motifs (FDR<0.05) were associated with their proximal target gene (TG) and represented as two nodes and one TF-TG edge. In total, there were 3,295,212-5,900,174 predicted TF-TG edges (FDR<0.05) across the five species (Mehta et al. 2021). This was encoded into a matrix of 1,131,812 predicted TF-TG edges (FDR<0.05), where each edge is present in at least two species (Mehta et al. 2021). To enable accurate analysis of GRN rewiring and retain relevant TF-TG interactions, all collated edges were pruned to a total of 843,168 TF-TG edges (FDR<0.05) where 1) the edge is present in at least two species; 2) edges are not absent in any species due to node loss or mis-annotation; and 3) based on the presence of nodes in modules of co-expressed genes in our previous study (Mehta et al. 2021).

### Three-node motif generation

Three-node motifs in our study are defined as a miRNA feed-forward loop (miRNA-FFL), where a TF is predicted to regulate a TG and a miRNA is predicted to directly regulate either the TF or TG (Fig. 3a). Three-node motifs (TF:TG:miRNA) were encoded by merging all combinations of predicted TF and miRNA interactions of a TG.

### Three node motif analysis

For each species three-node motifs, all TF:miRNA nodes were extrapolated for all TGs and their frequency recorded (based on the same TF orthogroup and miRNA family classification). By reverse ranking the frequency of all TF:miRNA nodes in each species, the top 100 relationships were classified to test for any significant overlap of TFs and miRNAs in species-specific three-node motifs.

A presence-absence matrix of three-node motifs in each species was generated, and the number of TFBS and miRNA binding site gains and losses, against predictions in *O. niloticus*, were calculated for each species TG. The degree-corrected rewiring (*D_n_*) score of TF-TG interactions in each orthogroup, as inferred by the DyNet-2.0 package (Goenawan et al. 2016) implemented in Cytoscape-3.7.1 (Franz et al. 2015), was then mapped for GRN rewiring analysis.

### Hypergeometric tests for regulatory site gain and loss enrichment

The *phyper* function in R (v4.0.2) was used to test for enrichment of rewired (degree-corrected *D_n_* score >0.17) or low to non-rewired (degree-corrected *D_n_* score ≤ 0.17) genes in each of the eight models of TFBS and/or miRNA binding site gains, losses or no change. The *D_n_* score threshold of 0.17 (for rewired vs low to non-rewired) was set based on the mean *D_n_* score for all orthogroups and as a measure of significantly rewired genes based on our previous study (Mehta et al. 2021). A threshold of *p*-value <0.05 was used as a measure of significant enrichment.

### Calculating substitution rate at regulatory regions

To identify loci evolving at a faster rate than that expected under a neutral model, we used phyloP (Pollard et al. 2010) from the Phylogenetic Analysis with Space/Time Models (PHAST) v1.5 package (Hubisz et al. 2011). This was carried out on the previously published 5-way *multiz* multiple alignment file (MAF) centred on *O. niloticus* v1.1 (Brawand et al. 2014). An *O. niloticus* centred MAF was used as a reference owing to its phylogenetic position as an outgroup to study substitution rates within the cichlid phylogeny and radiating lake species. Using the *O. niloticus* centred MAF, a neutral substitution model was constructed using the previously published five cichlid phylogeny (Brawand et al. 2014) in phyloFit from PHAST v1.5 (Hubisz et al. 2011) by fitting a time reversible substitution ‘REV’ model. The multiple alignment was split by chromosome/scaffold and phyloP (Pollard et al. 2010) ran using the likelihood ratio test (LRT) and the ‘all branches’ test to predict conservation-acceleration (CONACC) scores for each site in the five species multiple alignment.

To obtain pairwise phyloP scores, we 1) created MAFs that are centric to each of the other four species by reordering the original *O. niloticus* centred MAF using mafOrder from UCSC kent tools v333. Regardless of the species, the same alignment information is therefore retained throughout the workflow; 2) removed all alignments that excluded the reference species using mafFilter from UCSC kent tools v333; 3) created sorted MAFs for all pairwise species combinations using the mafFilter function in mafTools v0.1 (Earl et al. 2014); 4) constructed a neutral substitution model for each pairwise combination using phyloFit from PHAST v1.5 (Hubisz et al. 2011) by fitting a time reversible substitution ‘REV’ model; 5) split each pairwise MAF by chromosome/scaffold; and 6) calculated substitution rates in phyloP (Pollard et al. 2010) using the likelihood ratio test (LRT) and the ‘all branches’ test to predict conservation-acceleration (CONACC) using each corresponding pairwise neutral substitution model. To compare CONACC scores of regulatory regions to neutrally evolving regions, fourfold degenerate sites were extracted from each genome using an in-house perl script that takes a gene annotation as gene transfer format (GTF) file, whole genome FASTA file and fourfold degenerate codon table as input. The phyloP scores were then mapped to fourfold degenerate sites and the four regulatory regions (3’ UTR excluding miRNA binding sites, up to 5kb gene promoter excluding TFBSs, 3’ UTR miRNA binding sites and up to 5kb gene promoter TFBSs) of each species using *bedtools-2.25.0* intersect (Quinlan and Hall 2010).

### Identification of pairwise variation between the five species

After stage three of the above where pairwise species MAFs are created by sorting and filtering each species centric MAF, there were a total of 20 MAFs representing all pairwise species combinations (5 ref species × 4 comparison species). Each pairwise species MAF was used to identify pairwise variation using a custom python script ‘*get_subs_from_maf.py*’. Pairwise (single-nucleotide) variants were mapped to the phyloP scores of four regulatory regions (3’ UTR, up to 5kb gene promoter, miRNA binding sites and TFBSs) using *bedtools-2.25.0* intersect (Quinlan and Hall 2010).

### Testing the significance of difference in conservation-acceleration (CONACC) scores of pairwise variation in regulatory regions

The significance of conservation-acceleration (CONACC) scores of pairwise polymorphic nucleotide sites in regulatory regions of all species was tested both within and between species using the *Wilcoxon* rank sum test. The *p-*values were adjusted for multiple test correction using the Benjamini-Hochberg (BH) method (Benjamini and Hochberg 1995). The adjusted *p-*value was recorded as either having a significant (adjusted *p-*value <0.05) or insignificant (adjusted *p-*value >0.05) difference in CONACC scores of pairwise polymorphic nucleotide sites in within and between regulatory regions of all five species.

### Testing the significance of conservation-acceleration (CONACC) scores in regulatory regions of adaptive trait genes

The significance of conservation-acceleration (CONACC) scores of pairwise polymorphic nucleotide sites in regulatory regions of 90 adaptive trait genes was tested by: 1) Randomly picking up to 90 no to low rewired genes (*D_n_* score ≤0.17) from our previous study (Mehta et al. 2021), 1000 times; and 2) testing (*Wilcoxon* rank sum) the difference in CONACC scores of pairwise polymorphic nucleotide sites in each regulatory region of the 90 random genes to the corresponding regulatory region of all 90 adaptive trait genes. The *p-*values were adjusted for multiple test correction using the Benjamini-Hochberg (BH) method (Benjamini and Hochberg 1995). The number of times (from 1000 tests) was recorded as either having a significant (Wilcoxon rank sum test, adjusted *p-value* <0.05) or insignificant (Wilcoxon rank sum test, adjusted *p-value* >0.05) difference in CONACC scores of pairwise polymorphic nucleotide sites in each regulatory region of all five species. The adjusted *p-*values derived from Wilcoxon rank sum tests, between CONACC scores of polymorphic nucleotide sites in the regulatory region of 90 adaptive trait genes were ranked, and reverse sorted, to identify significant (adjusted *p-value* <0.05) regulatory binding site turnover.

### Identification of segregating variants within binding sites

Pairwise variants of *M. zebra* were overlapped with single nucleotide polymorphisms (SNPs) in Lake Malawi species (Malinsky et al. 2018) using *bedtools-2.25.0* intersect (Quinlan and Hall 2010). The pairwise variants overlapping binding sites and lake species SNPs were then filtered based on the presence of the same pairwise variant in orthologous 3’ UTR alignments. This ensured concordance of whole-genome alignment derived variants with variation in 3’ UTR alignments and predicted binding sites. At each step, complementation of reference and alternative alleles was accounted for to ensure correct overlap. This analysis was not carried out to distinguish population differentiation due to genetic structure, but to instead map 3’ UTR regulatory variants onto a number of radiating cichlid species to link to phylogenetic and ecological traits.

### Phylogenetic independent contrasts

Phylogenetic independent contrasts (PICs) were carried out to statistically test the effect of fitting the 73 Lake Malawi species phylogeny (Malinsky et al. 2018) to the covariance of segregating TFBSs and miRNA binding sites, visual (wavelength palette) and ecological traits (habitat and foraging habit/diet). This involved 1) categorically coding the genotypes of segregating regulatory sites, visual trait and ecological measurements for each of the 73 Lake Malawi species (119 individuals), and 2) using the *ape* package (v5.4.1) in R (v4.0.2) to apply the PICs test (Felsenstein 1985) on all correlations with the binding site genotype (genotype vs wavelength palette, genotype vs habitat, and genotype vs foraging habit/diet). PICs assumes a linear relationship and a process of Brownian motion between traits, and thus, for each combination of data, scatterplots were first generated. To test for any change in the correlation owing to phylogenetic signal, the regression model was compared between the relationships both excluding and including the Lake Malawi phylogeny (Malinsky et al. 2018).

## Declarations

### Data Availability

Data underlying this article is available in the article and in its online supplementary material.

### Consent for publication

Not applicable.

### Competing interests

The authors declare that they have no competing interests.

## Acknowledgments

We thank the BROAD institute and the Cichlid Genome Consortium for providing full access to genomic data.

## Funding

This work was supported by the Biotechnological and Biosciences Research Council (BBSRC), part of UK Research and Innovation, Institute Strategic Programme (grant number BB/J004669/), Core Strategic Programme Grants (grant numbers BB/P016774/1, BBS/E/T/000PR9817, BB/CSP17270/1, BB/CCG1720/1) to TKM, LPD, WN, FDP, and WH at the Earlham Institute, and acknowledge the work delivered via the Scientific Computing group, as well as support for the physical HPC infrastructure and data centre delivered via the NBI Computing infrastructure for Science (CiS) group. This work was also supported by a National Science Foundation (NSF) career award (grant number 1350677 to SR) and the McDonnell foundation at The Wisconsin Institute for Discovery.

## Authors’ contributions

TKM ran gene ontology (GO) enrichment, three-node motif generation, analysed three-node motifs, tested regulatory site gain and loss, calculated substitution rates, identified pairwise variants, tested the significance of CONACC scores, identified segregating binding sites, and ran phylogenetic independent contrast (PICs) tests; LPD ran miRNA target prediction; WN ran TF motif scanning; TKM and WH wrote the manuscript with input from LPD, WN, SR and FDP.

## Authors’ information

Twitter handles: @TK_mehta (Tarang K. Mehta); @LucaPensoDolfin (Luca Penso-Dolfin); @nashalselection (Will Nash); @sroyyors (Sushmita Roy); @ScienceisGlobal (Federica Di-Palma); @WHaerty (Wilfried Haerty).

## Corresponding authors

Correspondence to

## Notes

### Competing Interest Statement

The authors have declared no competing interest.

### Summary of Updates

Main and supplementary text have been updated with additional analyses.

