## Supplementary Text for "Evolution of miRNA binding sites and regulatory networks in cichlids"

**Supplementary Information**

**Results**

**miRNA binding site prediction in 3’ UTRs of genes**

Using a filtered set of 15,390,993 binding sites, unique predicted miRNA binding sites of a target gene (TG) 3’ UTR were classified as miRNA:TG edges in each species. These edges were used to compare the total number of common and unique miRNA binding sites across all orthologous TGs based on miRNA:TG overlap along the phylogeny (Fig. 1b). In Fig. 1b, the numbers at ancestral nodes represent unique binding sites that could have existed at the common ancestor for those species. As an example, the miR-153:dlx5a edge is present in all five species and therefore counted at the Anc4 node as a common binding site for all five species. However, the miR-33:twist1b edge is only present in *M. zebra, P. nyererei* and *A. burtoni,* and therefore counted at the Anc2 node, common only to the haplochromine species. The miR-33:twist1b edge therefore represents a binding site that was likely gained in the common ancestor for all haplochromines since it is not found in either *O. niloticus* or *N. brichardi*. Otherwise, if an edge e.g. miR-126:trim65 is absent in an intermediate species e.g. *P. nyererei* but present in some other species e.g. *M. zebra* and *A. burtoni* then we count it at the ancestral node e.g. Anc2 of the deepest lying species e.g. *A. burtoni.* These patterns of binding site gain and loss occur along the phylogeny; there appears to be a significant loss of binding sites from Anc4 to Anc3, but then a gain of binding sites (from Anc3) in the haplochromine common ancestor (Anc2), and then a loss in Anc1 (from Anc2) (Fig. 1b). The gain in Anc2 coincides with the rapid expansion of haplochromines, and therefore suggests an important regulatory role in haplochromine diversification, that could have become simplified i.e. purifying selection, in *M. zebra* (Lake Malawi) and *P.nyererei* (Lake Victoria).

Using the 19,613,903 miRNA binding sites in annotated 3’ UTRs of 21,871 orthogroups across five cichlid species (Fig. 1a), we predicted an average of 179 (*M. zebra* – 186; *P. nyererei* – 147; *A. burtoni* – 200; *N. brichardi* – 138; and *O. niloticus* – 225) miRNA binding sites per 3’ UTR. After filtering for 18,799 co-expressed orthogroups, we retained 15,390,993 predicted binding sites across the five species (Fig. 1a, Supplementary Fig. S1). This retained an average of 218 (*M. zebra* – 218; *P. nyererei* – 179; *A. burtoni* – 241; *N. brichardi* – 182; and *O. niloticus* – 272) miRNA binding sites per 3’ UTR. By comparing the number of common and unique binding sites across orthologous 3’ UTR sequences (Fig. 1b), we identified 31,186 (*P. nyererei*) to 128,831 (*A. burtoni*) unique binding sites in the five species (Fig. 1b). These findings are not biased by genome completeness or annotation quality, since *N. brichardi* has the fewest annotated protein-coding genes (20,199 – compared to 20,611-24,559 in the other four species) and lowest genome contig N50 (13.2 kb – compared to 20-29.3 kb in the other four species), whereas *O. niloticus* has the most complete genome and highest protein coding gene count (Brawand et al. 2014).

Whilst there is a similar representation based on the 10 topmost miRNA families and their number of targets, there is variability in the number of target genes per family across the five species (Supplementary Fig. S2). For example, *miR-737* is unique and has the second most binding sites in *A. burtoni* and *miR-15c* binding sites are under-represented in *N. brichardi* (Supplementary Fig. S2). We performed further analysis to test the effect of the *miR-15c* seed mutation (AGCAGCG) on the number of predicted miR-15c binding sites in *N. brichardi* compared to the other species. Since Targetscan7 uses seed matching as one of its predictive features (Agarwal et al. 2015), we re-ran Targetscan7 (same approach as described in the *Materials and Methods*) using the mutated ‘AGCAGCG’ seed sequence of *miR-15c* to predict binding sites in 3’ UTR sequences of all species, and then compared the total number of binding sites outputted. If there is indeed an effect of the seed mutation on the number of predicted *miR-15c* binding sites in *N. brichardi*, then we would expect fewer predicted binding sites in *N. brichardi* compared to the other species. If there is no effect, then we would expect more *miR-15c* binding sites in *N. brichardi* compared to the other species, as the mutated seed has instead affected the number of predicted sites in the other species. Using the mutated seed sequence, we found that the fewest *miR-15c* binding sites were still predicted in *N. brichardi* (6711 binding sites) compared to the other four species (7766-11047 binding sites). These results therefore show that mutations of the *miR-15c* seed sequence are likely to be associated with under-represented *miR-15c* binding sites in *N. brichardi* compared to the other four species.

Finally, gene ontology (GO) enrichment of target genes for the 10 topmost miRNA families highlight terms that are both common e.g. membrane and signal transduction, and unique e.g. ATP binding and zinc ion binding (FDR<0.05) between the five species (Supplementary Fig. S3). Genes associated with similar functions have been previously implicated in miRNA/gene target pairs with negatively correlated expression in Midas cichlids (Franchini et al. 2019), and miRNA expression in B chromosomes of *A. latifasciata* (Nascimento-Oliveira et al. 2021)*.*

**Differential miRNA binding site usage highlights rewiring at the post-transcriptional level**

We predicted functional divergence of binding sites based on the identification of more than one miRNA family that positionally overlap in a gene’s 3’ UTR (Supplementary Fig. S4). For example, at position 858-865 in the 3’ UTR alignment of the visual opsin gene, *sws2a*, the binding site for miR-725 is conserved between *A. burtoni* and *O. niloticus* (Supplementary Fig. S4a) however, at position 564-571 in the same alignment, the binding site for miR-142 is predicted in *A. burtoni* but miR-1388 in *O. niloticus* (Supplementary Fig. S4b), and therefore functionally diverged due to discrete nucleotide variations. This method was used to predict functional divergence and allowed for us to assess the number of shared sites utilised by either the same e.g. miR-725 (Fig. 2a, Supplementary Fig. S5a) or different e.g. miR-142 and miR-1388 (Fig. 2b, Supplementary Fig. S5b) miRNA families between species. In total, 50,212 sites are conserved across all species (Anc4 node, Fig. 2a). This includes miRNA binding sites in 3’ UTRs of genes that have been previously implicated with cichlid adaptive traits like, for example, the jaw development gene *dlx5* (Bloomquist et al. 2017) with three miRNA targets e.g. miR-153, and a gene associated with deep-water adaptation, *zmiz* (Hahn et al. 2017) with 49 targets e.g. miR-17. The haplochromine species share the second most number (Anc2 node: 32,087) of binding sites (Fig. 2a). An example of a shared site in the haplochromines includes *rxrg*:miR-722; the target gene, *rxrg*, is a nuclear hormone receptor that promotes the differentiation of long-wave sensitive (LWS) cones and supresses *sws1* cone opsin expression in mice (Roberts et al. 2005), and miR-722 has been linked with LWS opsin regulation in cichlids (O'Quin et al. 2011). The highest number of pairwise shared sites of same miRNA families are found between *M. zebra* and *A. burtoni* (42,936) and overall, in all pairwise comparisons with *O. niloticus* (21,944-39,907 sites) (Supplementary Fig. S5a). An example of a shared site between *M. zebra* and *A. burtoni* includes *tfap2c*:miR-726; the target gene, *tfap2c*, is a transcription factor (TF) that has been previously implicated in zebrafish retinal-pigmented epithelium (RPE) programs (Buono et al. 2021), and miR-726 is expressed in zebrafish retina (Kloosterman et al. 2006) and implicated in cichlid opsin gene regulation (O'Quin et al. 2011). The characterisation of shared sites therefore identifies predicted miRNA regulation of genes associated with traits of cichlid phenotypic diversity, such as the visual system (Carleton 2009).

Counter to this, there is more miRNA family divergence within the haplochromine lineage (Anc2:3163 and Anc1:3200 shared sites) compared to all other phylogenetic comparisons (Anc4:1 and Anc3:17 shared sites) (Fig. 2b). Accordingly, the highest number of pairwise shared sites utilised by different miRNA families is observed in the haplochromine lineage, between *A. burtoni* and *M. zebra* (6,972 sites) (Supplementary Fig. S5b). The lowest number of pairwise shared sites is observed between two of the most evolutionary diverged species, *O. niloticus* and *P. nyererei* (1,108 sites) (Supplementary Fig. S5b). Many more instances of rewired binding sites (3,163) are therefore utilised by different miRNA families between the haplochromine species and thus, less prone to reuse binding sites that are only present in basal representatives (2,083 sites in *N. brichardi* and *O. niloticus*) (Supplementary Fig. S5b).

#### Comparative analysis of three-node motifs identifies increased novel network architecture between the five cichlid species

We previously generated transcription factor (TF) - target gene (TG) networks for the five cichlid species (Mehta et al. 2021). Since a GRN can be composed of both transcriptional activation and repression, we develop upon our previous findings (Mehta et al. 2021) by instead focusing on ‘three-node motifs’ (Alon 2007) as a measure of network architecture and evolutionary constraint, and more specifically, a topology representative of a miRNA feed-forward loop (miRNA-FFL) (Fig. 3a). According to the miRNA-FFL model (Fig. 3a), we started with 37,320,950 three-node motif edges present in any of the five species (Supplementary Fig. S6). To avoid any bias of gene loss or mis-annotations in motif comparisons across all species, we first focused on 17,987,294 three-node motif edges composed of 1-to-1 orthologous nodes (of TFs and TGs) across the five species (Supplementary Fig. S7). In this set, only 467,279 (3%) three-node motif edges are conserved across all five species; 155,984 (1%) – 380,748 (2%) edges are conserved in all four species combinations; 420,942 (2%) edges are conserved in the haplochromines only; and 173,750 (1%) – 641,715 (4%) edges are conserved in all pairwise combinations (Supplementary Fig. S8, Table S2). Instead, 1,321,875 (7%) – 3,124,263 (17%) three-node motifs are unique to each species (Supplementary Fig. S8, Table S2). For the zebrafish housekeeping gene, *ef1a* (Casadei et al. 2011), we find 79 three-node motifs conserved in all five species, but consistent with phylogenetic relatedness, we find more (111) three-node motifs conserved in the haplochromines only. Compared to the number of novel three-node motifs in each species, the lower level of conserved three-node motifs highlights binding site turnover and rewiring between the five species. However, to determine whether this is due to either TF or miRNA binding site turnover, the three-node motifs also need to be composed of 1-to-1 orthologous nodes (of miRNAs and TGs) across the five species too. This was done by classifying binding sites of each miRNA into families; for example, binding sites of miR-27a/miR-27b/miR-27c become a single orthologous group known as miR-27, and if there are binding sites of any member present in each of the five species, then it is deemed 1-to-1 orthologous. Taking the 992 (On-198, Nb-183, Ab-243, Mz-185, Pn-183) cichlid miRNA mature sequences used for target prediction, a total of 68/172 (40%) miRNA families are classified as non 1-to-1 orthologous miRNAs. Upon filtering the three-node motifs for these miRNAs, a total of 17,987,294/21,708,657 (83%) three-node motif edges remained, composed of 1-to-1 orthologous nodes (of TFs, miRNAs and TGs) across the five species (Supplementary Fig. S6).

Based on three-node motifs across the five species, the pattern of increased novel/species-specific network architecture is also observed in the differing patterns and frequency of 115,031 unique TF:miRNA relationships (see *Materials and Methods*). In total, the frequency of 115,031 unique co-occurring TF:miRNA relationships ranges from 1 to 2143 in 1-to-1 orthologous three-node motifs across the five species. A total of 2,222 (2%) TF:miRNA relationships are conserved in the haplochromines including, for instance, a relationship between *miR-20a* and GATA2A, a TF implicated in GRN rewiring of the dim-light vision gene, *rhodopsin (rho)* (Mehta et al. 2021). In all five species, the top 100 most frequent TF:miRNA relationships in 1-to-1 orthologous node three-node motifs co-occur a total of ≥1069-2143 times, identifying a total 255 unique TF:miRNA relationships. Only 10/255 (4%) e.g. EGR2B:*miR-129* are conserved across all five species; 12/255 (5%) e.g. IRF7:*miR-23a* are conserved in the haplochromines; and 123/255 (48%) e.g. *O. niloticus* GATA2A:*miR-23* are unique to any one species (Supplementary Fig. S10). Using a set of 90 candidate teleost and cichlid trait genes associated with phenotypic diversity from previous studies (Supplementary Table S3), we note that the highest frequency of TFs in TF:miRNA relationships in all five species are of brain development and neurogenesis associated TFs (Bloomquist et al. 2017) e.g. NEUROD1:miR-129 (360-452 occurrences), NEUROD2:miR-23a (173-226 occurrences), and GATA6:miR-101a (93-136 occurrences). In total, 2/90 genes are targets in three-node motifs of at least one of the top 100 most frequent TF:miRNA relationships, and have relationships conserved across all five species; including the homeobox pathway gene (Bloomquist et al. 2017), *dlx5a* e.g. MXI1:miR-129 (1381-1925 occurrences) and the photoreceptor development gene (Hahn et al. 2017), *sgpp1* e.g. EGR2:miR-129 (1493-1899 occurrences) (Supplementary Fig. S10). Whereas, 10/90 genes are targets in three node motifs with at least one of the top 100 most frequent TF:miRNA relationships, and as an example, either have 1) relationships conserved in two to four species, including the homeobox pathway gene (Bloomquist et al. 2017), *nkx2-1* e.g. ZNF384L:miR-23a (1314-1786 occurrences), and the notch pathway gene (Bloomquist et al. 2017), *notch3* e.g. MXI1:miR-2187b (1171-1649 occurrences); or 2) species-specific relationships, including the fibroblast growth factor pathway gene (Bloomquist et al. 2017), *fgfr1* e.g. RELB:miR-20009 (2583 occurrences in *A. burtoni*) and the morphogenesis gene also found to be fast evolving in cichlids (Brawand et al. 2014), *bmpr1* e.g. IRF7:miR-27b (1448 occurrences in *M. zebra*) (Supplementary Fig. S10).

We also tested whether any of the 90 adaptive trait genes are enriched as species-specific targets genes (TGs) to make associations to any lineage-specific phenotypes. We tested this by 1) extracting the TGs of TF:miRNA relationships that are unique to each species; and then 2) running a hypergeometric enrichment test of each set of species-specific TGs against the 90 adaptive trait genes (from Supplementary Table S3) in the corresponding species. However, we found no significant (*p*-values: 0.42 - 0.67) enrichment of adaptive trait genes that are targets for species-specific TF:miRNA relationships. This is to be expected as we only tested 90 cichlid adaptive trait genes against an average of 5658 target genes across the five species. Nonetheless, the TGs we identify of species-specific TF:miRNA relationships could represent genes associated with lineage specific adaptive traits however, this requires further testing using a wider set of cichlid adaptive traits genes. Overall, we find that 23-27 (an average of 25) out of the 90 adaptive trait genes are found to be targeted by the 5191-6092 (0.4-0.5%) unique species-specific TF:miRNA relationships. This includes, as examples: 1) *chl1,* associated with cichlid neural adhesion (Bloomquist et al. 2017), that is a species-specific target in both *M. zebra* and *P. nyererei* of only 3 co-regulatory relationships e.g. TBX21:*miR-30a*; and 2) *draxin,* a gene that could be associated with cichlid eye morphogenesis (Hahn et al. 2017), is targeted in all five species by 266 co-regulatory relationships e.g. EGR1:*miR-214*.

Overall, by focusing on three-node motifs that can, based on associations to previous literature, be better functionally associated than single nodes alone e.g. cichlid miRNAs, we may be able to highlight network rewiring events associated with particular traits of cichlid phenotypic diversity.

**Network rewiring is associated with different models of regulatory binding site turnover in three-node motif across species**

To determine the relative impact of TFBS/miRNA gain and loss in three-node motifs towards GRN rewiring, we implemented further models of TFBS and/or miRNA binding site evolution. Considering the previous analysis (Fig. 3b) can include overlapping models of binding site evolution for each TG, we implemented a total of eight models of TFBS and/or miRNA binding site evolution, including ‘no change’, using *O. niloticus* as a reference. This allowed us to determine whether any of the eight models of binding site evolution could be significantly associated (hypergeometric *p*-val <0.05) to any of the previously described rewired 1-to-1 orthologs (Mehta et al. 2021). Using the previously described (Mehta et al. 2021) mean degree-corrected DyNet rewiring (*D_n_*) score (Goenawan et al. 2016) for all 6,802 1-to-1 orthogroups, a total of 6,542 orthogroups were assigned as significantly rewired (degree-corrected *D_n_* score >0.17) and 260 orthogroups as low to non-rewired (degree-corrected *D_n_* score ≤0.17) (see *Materials and Methods*). Since rewiring was measured in our previous study based on TFBS divergence (Mehta et al. 2021), we expect that TFBS gain/loss, instead of miRNA binding site gain/loss, to have the largest effect on significantly rewired (degree-corrected *D_n_* score >0.17) orthologs. Overall, a higher number of orthogroups are rewired (than low to non-rewired) under all eight models of binding site evolution (Fig. 4a, Supplementary Fig. S11, Table S5). Under all eight models, very few (6-10%) of the 6,802 1-to-1 orthogroups exhibit any of the four models with ‘no change’ in binding sites, whereas 71% and 49% are rewired (mean *D_n_* score = 0.21) and exhibit either ‘TFBS and miRNA binding site loss’ or ‘TFBS gain and miRNA binding site loss’ in the four species compared to *O. niloticus* (Fig. 4a, Supplementary Fig. S11, Table S5). After running hypergeometric tests to assess over-representation in each species, the rewired 1-to-1 orthologs (*D_n_* score >0.17) are most associated with TFBS and miRNA binding site gain in *M. zebra* (*p*-value =0.03) and *N. brichardi* (*p*-value =0.0007), TFBS gain and miRNA binding site loss in *P. nyererei* (*p*-value =0.001), and TFBS loss and miRNA binding site gain in *A. burtoni* (*p*-value =0.009) (Supplementary Table S6). As expected, this suggests that TFBS-based rewiring of GRNs from our previous study (Mehta et al. 2021), was largely impacted by TFBS gain along the phylogeny. This is therefore indicative of a discrete impact of GRN rewiring based on miRNA binding site loss. Overall, these results suggest that different models of regulatory binding site evolution have impacted GRN rewiring in the studied cichlid lineages.

**Regulatory binding site turnover in three-node motifs is associated with network rewiring of adaptive trait genes**

We previously showed that out of 31 1-to-1 orthologous adaptive trait genes, nine e.g. *rh2* and *draxin,* have comparatively higher rewired networks (*D_n_* score 0.17 ± 0.03 SD) than all 1-to-1 orthologs (Mehta et al. 2021). Of the 31 1-to-1 orthologous genes implicated in teleost and cichlid phenotypic diversity (Supplementary Table S3), 2/31 e.g. *dlx5* have the same and 29/31 e.g. *draxin* have different models of binding site evolution in three-node motifs between the four species (compared to *O. niloticus* as a reference). For example, the homeobox pathway gene (Bloomquist et al. 2017), *dlx5,* and neurogenesis gene (Bloomquist et al. 2017), *chl1*, have TFBS and miRNA binding site gain in all four species (Supplementary Fig. S12). On the other hand, *sws2a* (short-wave-sensitive) opsin, utilised as part of the long-wavelength palette in *P. nyererei* and *A. burtoni*, has TFBS gain and miRNA binding site loss in *P. nyererei*, TFBS and miRNA binding site gain in *A. burtoni*, but TFBS and miRNA binding site loss in the other two species (Fig. 4b). Further to this, *rh2b* (middle-wave-sensitive) opsin, utilised as part of the short-wavelength palette in *M. zebra* and *N. brichardi*, has TFBS loss and no change in miRNA binding site in *M. zebra*, but TFBS and miRNA binding site loss in the other three species (Fig. 4b). Similarly, the brain development and neural differentiation gene, *neurod2,* and the fast evolving morphogenesis gene, *bmpr1b* (Brawand et al. 2014), which are predicted to be regulated by two novel miRNAs (miR-10032: *neurod2* and miR-10029: *bmpr1b*) in cichlids (Brawand et al. 2014), are both rewired (*D_n_* score >0.17) and have different models of binding site evolution between the four species. Compared to *O. niloticus* as a reference*, neurod2* has no change in TFBS and miRNA binding site loss in *M. zebra*, TFBS and miRNA binding site loss in *P. nyererei* and *N. brichardi,* and TFBS gain and miRNA binding site loss in *A. burtoni.* Instead, *bmpr1b* has TFBS gain and miRNA binding site loss in *M. zebra*, no change in TFBS and miRNA binding site loss in *P. nyererei*, and TFBS and miRNA binding site loss in *A. burtoni* and *N. brichardi* (Fig. 4b). On the other hand, a closely related proneural gene, *neurod1,* which is also rewired (*D_n_* score >0.17), has two different models of binding site evolution between species; having TFBS loss and miRNA binding site gain in *M. zebra* and *A. burtoni*, but TFBS and miRNA binding site loss in *P. nyererei* and *N. brichardi* (Fig. 4b). Other examples of which include the organogenesis fibroblast growth factor pathway gene (Bloomquist et al. 2017), *fgfr1* (*D_n_* score = 0.23), that has no change in TFBS and miRNA binding site loss in *M. zebra*, TFBS loss and miRNA binding site gain in *A. burtoni*, but TFBS and miRNA binding site loss in the other two species; the neurogenesis gene (Bloomquist et al. 2017), *nrp1a* (*D_n_* score = 0.23), that has TFBS and miRNA binding site gain in *P. nyererei,* TFBS loss and miRNA binding site gain in *A. burtoni*, but TFBS and miRNA binding site loss in the other two species (Fig. 4b); the homeobox pathway gene (Bloomquist et al. 2017), *dlx3b* (*D_n_* score = 0.19)*,* has TFBS gain and miRNA binding site loss in *M. zebra*, TFBS and miRNA binding site gain in *P. nyererei,* TFBS loss and miRNA binding site gain in *A. burtoni*, and TFBS and miRNA binding site loss in *N. brichardi* (Fig. 4b). Differing models of binding site evolution are also observed for low to non-rewired (*D_n_* score ≤0.17) 1-to-1 orthologs, including the photoreceptor development gene (Hahn et al. 2017), *sgpp1* (*D_n_* score = 0.15), which has TFBS and miRNA binding site gain in *M. zebra* and *P. nyererei*, but TFBS gain and miRNA binding site loss in the other two species (Fig. 4b).

Owing to the large phenotypic differences between the five species, as well as tissue-specific gene expression and TF regulatory differences (Mehta et al. 2021), we expect that genes associated with such phenotypic traits e.g. bone morphogenetic proteins (BMPs) in jaw shape/function (Albertson and Kocher 2006) and fibroblast growth factors (FGFs) in organogenesis (Popovici et al. 2005; Thisse and Thisse 2005), are likely regulated by differential patterns of TF and miRNA regulatory divergence.

**Discrete changes at regulatory sites are associated with binding site turnover in three-node motifs**

We studied TF and miRNA regulatory divergence by assessing the rate of evolution in regulatory regions and fourfold degenerate sites (as a proxy for neutrally evolving sites) compared to a neutral model. We compared the accelerated scores between all five genomic regions (fourfold degenerate, 3’ UTR, up to 5kb gene promoter, miRNA binding sites and TFBSs), either within or between species. With the exception of accelerated scores in 3’ UTR vs miRNA binding sites in *P. nyererei* and 5kb gene promoter vs miRNA binding sites in *N. brichardi-M. zebra* and *N. brichardi-A. burtoni* pairwise comparisons (adjusted *p-value* =0.1), there is a significant difference (Wilcoxon rank sum test, adjusted *p-value* <0.05) in the proportion of accelerated sites between all five regions both within and between species (Supplementary Table S11).

Since our previous study found that discrete regulatory mutations are able to drive GRN rewiring events (Mehta et al. 2021), we hypothesised that such mutations could account for some of the accelerated regulatory sites. Using pairwise polymorphic nucleotide sites in each of the four regulatory regions (Supplementary Table S12), we find that on average, 28% of accelerated sites in regulatory regions are driven by variation in a single species (Supplementary Fig. S15, Supplementary Table S13-14). With the exception of accelerated scores of single nucleotide variation in 5kb gene promoter vs miRNA binding sites in *P. nyererei* (adjusted *p-value* =0.5), there is a significant difference (Wilcoxon rank sum test, adjusted *p-value* <0.05) in the proportion of accelerated sites accounted for by variation in a single species between all four regulatory regions both within and between species (Supplementary Table S15).

**Discrete changes at regulatory sites are preferentially associated with regulatory binding site turnover in adaptive trait genes**

Based on our previous work, we found that a total of 60/90 adaptive trait genes had comparatively higher rewired regulatory networks (based on TFBSs) than all orthologs (Mehta et al. 2021). Here, we were interested in regulatory binding site turnover i.e. fast evolving pairwise polymorphisms, in regulatory regions and three-node motifs of all 90 adaptive trait genes (Supplementary Table S3). In these 90 genes and across all species comparisons, a total of 20,691 (3’ UTR), 110,584 (up to 5kb gene promoter), 1,850 (miRNA binding sites), and 31,268 (TFBSs) pairwise polymorphic sites are evolving at a faster rate than that expected under a neutral model. We test how many times this is significant compared to a random subset of all genes (see *Materials and Methods*); with the exception of *A. burtoni* 3’ UTR and miRNA binding sites in P. nyererei, *A. burtoni,* and *O. niloticus*, the majority (563-991) of 1000 Wilcoxon rank sum tests of CONACC scores of pairwise polymorphic sites in four regulatory regions, between 90 no to low rewired (*D_n_* score ≤0.17) and 90 adaptive traits genes, show a significant (adjusted *p-value* <0.05) difference in the proportion of accelerated sites (Supplementary Table S16). The most notable differences (691-991/1000 Wilcoxon rank sum tests, adjusted *p-value* <0.05) of which, are found in the proportion of accelerated sites in TFBSs of 90 adaptive trait gene promoter regions (Supplementary Table S16).

By reverse ranking significantly different (adjusted *p-value* <0.05) accelerated sites between all regulatory regions, we were able to identify 17 adaptive trait genes with significant turnover between TF and miRNA binding sites (Supplementary Table S17-18). In these 17 adaptive trait genes and across the three species comparisons, a total of 240 pairwise polymorphic nucleotide sites in miRNA binding sites (*M. zebra* - 83*, P. nyererei* - 47*,* and *O. niloticus* - 110) and 1,797 sites in TFBSs (*M. zebra* - 411*, P. nyererei* - 471*,* and *O. niloticus* - 915) are evolving at a significantly faster rate (adjusted *p-value* <0.05) than that expected under a neutral model (Supplementary Table S18).

**Discrete changes at regulatory regions of adaptive traits genes segregate according to phylogeny and ecology of radiating cichlids**

We finally studied regulatory binding site turnover in three-node motifs as a result of between species TFBS and miRNA binding site variation in the context of phylogeny and ecology of lake species. In the main text, we largely focus on genes (*sws1* and *rho)* associated with spectral tuning of visual systems (Fig. 5, Supplementary Fig. S16-19). However, we also note variation at accelerated regulatory sites of other adaptive trait genes in the Lake Malawi species, *M. zebra,* used as a reference here, to compare regulatory binding site turnover in three-node motifs that could be associated with the ecology of sequenced Lake Malawi species. If the TFBSs and miRNA binding sites are likely functional, we hypothesise that radiating species within the same clade would share conserved regulatory genotypes, to possibly regulate and perform same/similar functions; whereas distally related species would segregate at the corresponding regulatory site. We identified 94/494 sites with between species variation across 73 Lake Malawi species, that also exhibited flanking sequence conservation, representative of shared ancestral variation. Amongst these sites includes between species variation at a three-node motif for the jaw and pharynx developmental gene, *msx1b* (Bloomquist et al. 2017). Regulation of *msx1b* by a three-node motif consisting of DNAJC2, a transcriptional activator (Klisch et al. 2006), and miR-129, associated with osteoblast differentiation and bone formation (Yin et al. 2020), could regulate jaw bone development specifically in Lake Malawi species, but not *P. nyererei, A. burtoni* or *N. brichardi* (Supplementary Fig. S24-25). Phylogenetic independent contrast analysis (Felsenstein 1985) of the DNAJC2:*msx1b* genotypes against visual traits and ecology of each of the 73 Lake Malawi species, highlights very little change in correlation once the phylogeny is taken into account and a regression model fitted (Supplementary Fig. S26). However, due to a lack of sufficient characters, a regression model could not be fitted for miR-129 and therefore, PICs could not be tested (Supplementary Fig. S27).
