## Supplementary Figures for "Evolution of miRNA binding sites and regulatory networks in cichlids"

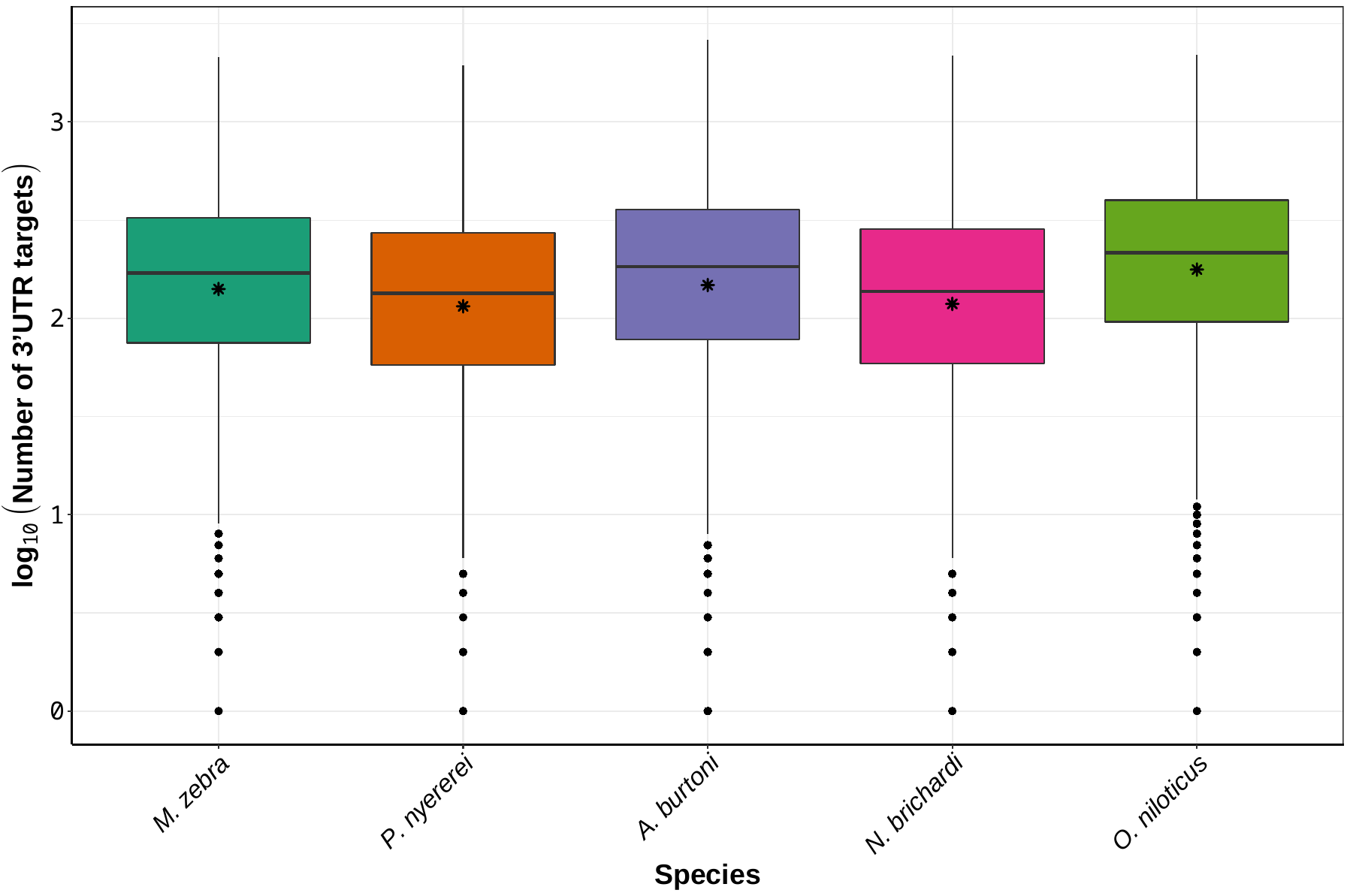

**Fig. S1 - Number of miRNA targets in 3’ UTR regions of each co-expressed gene in each species.** Box plot of log_10_ transformed counts of number of miRNA targets per co-expressed gene 3’ UTR in each species. Outliers are represented by external dots and mean values are shown as internal stars.

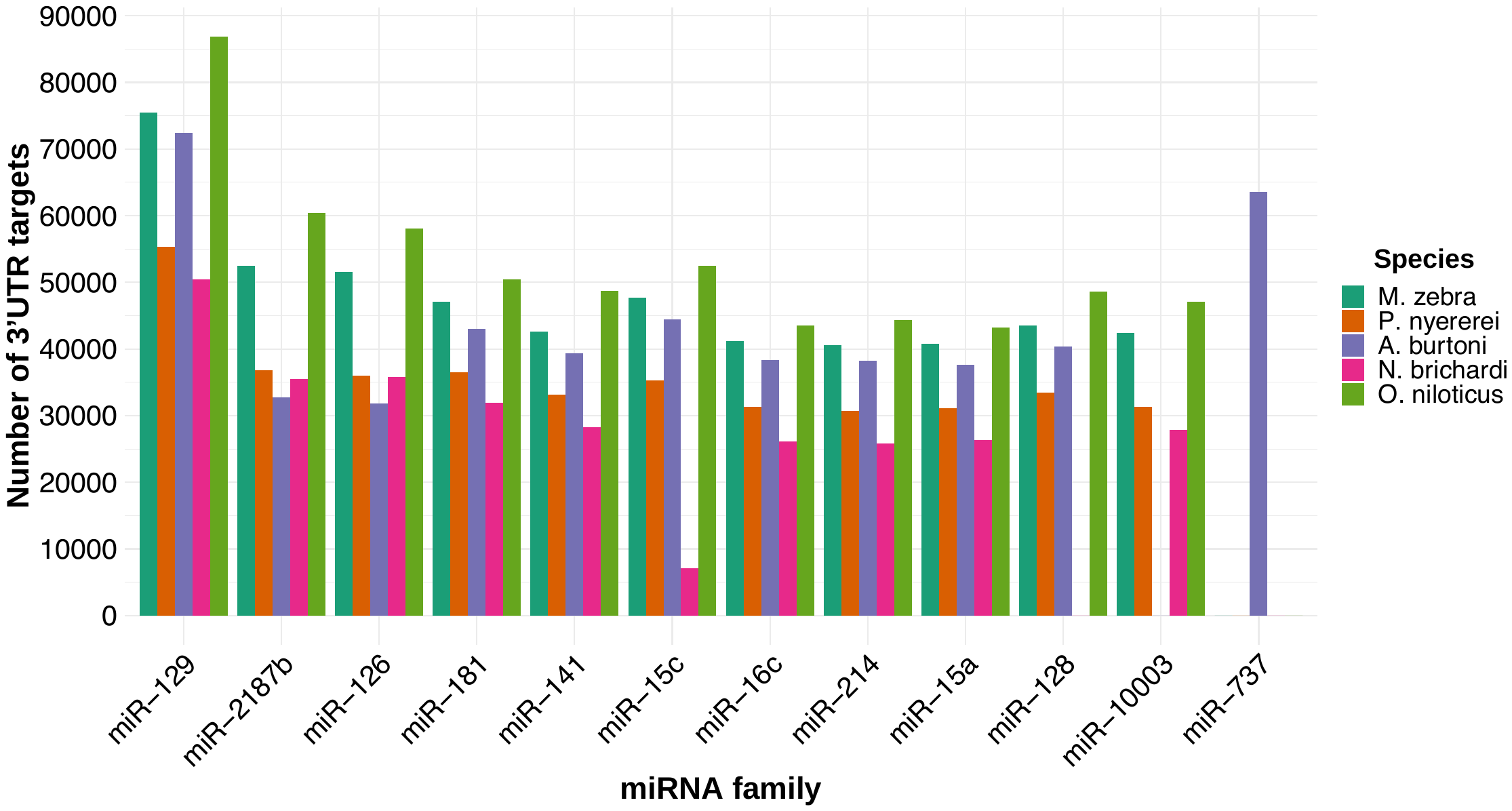

**Fig. S2 - Top 10 miRNA families targeting 3’ UTRs in each of the five cichlids.** Number of 3’ UTR targets (y-axis) and miRNA family (x-axis).

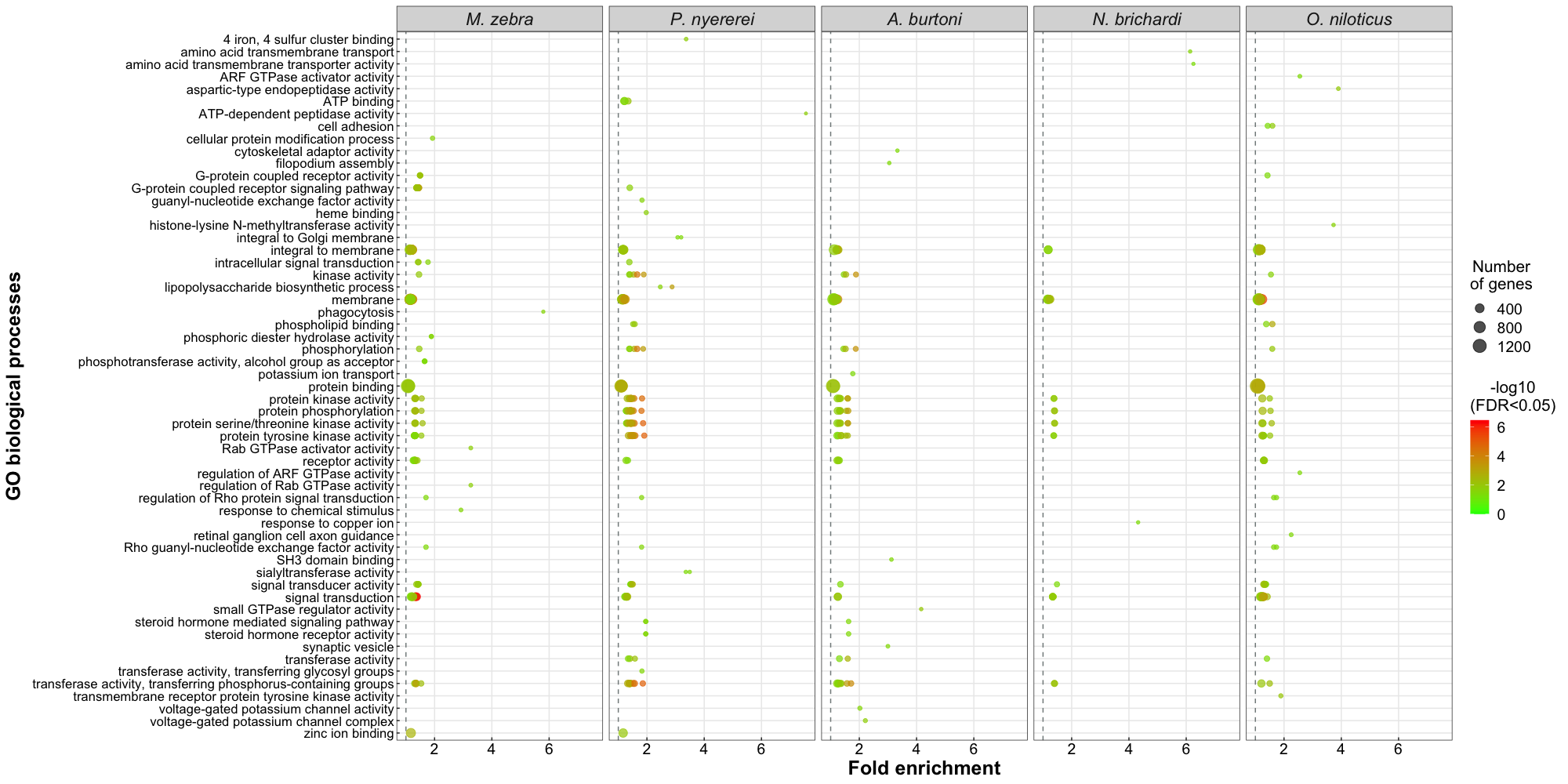

**Fig. S3 – Gene Ontology (GO) enrichment of top 10 miRNA families with 3’ UTR targets in each of the five cichlid species.** Circles show enriched biological processes (y-axis) of significance (*log_10_* FDR <0.05, heatmap to right) and *log_10_* fold enrichment (x-axis) values of miRNA families across all five species. Number of enriched genes for each term shown by size of each circle.

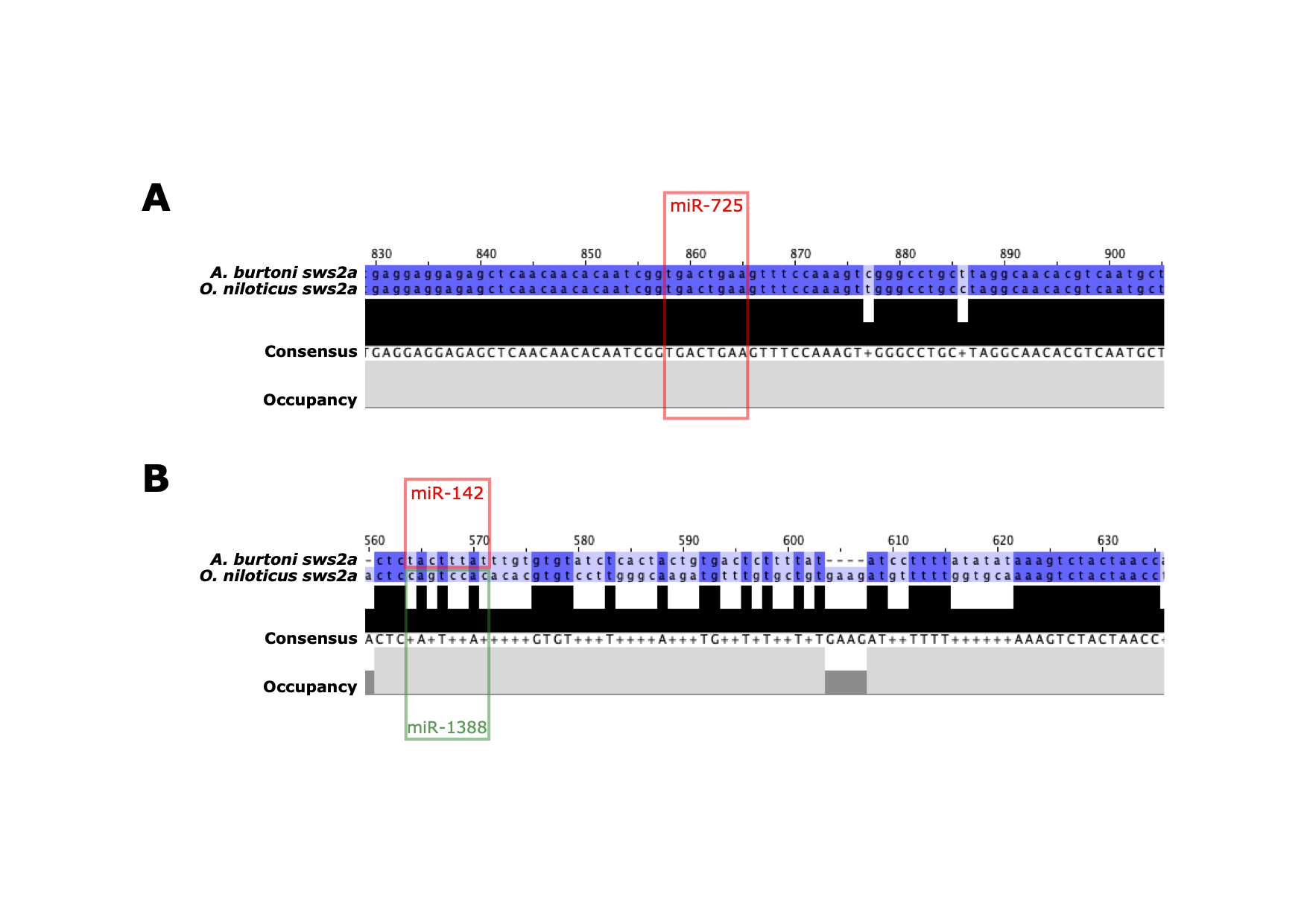

**Fig. S4 – Example of miRNA binding site conservation and divergence. (A)** Conservation of a miRNA binding site, miR-725 (red box), in the 3’ UTR of *A. burtoni* and *O. niloticus sws2a* gene*.* **(B)** Divergence of a miRNA binding site where at the same position in the alignment (full positional overlap), miR-1388 (green box) and miR-142 (red box) are predicted in the 3’ UTR of *A. burtoni* and *O. niloticus sws2a* genes respectively.

**Fig. S5 – Evolution of miRNA binding sites along the five cichlid phylogeny.** Number of shared and non-shared target sites based on miRNA binding site overlap in multiple 3’ UTR alignments are shown at ancestral nodes and branches for **(A)** same miRNA families and **(B)** different miRNA families with all pairwise comparisons.

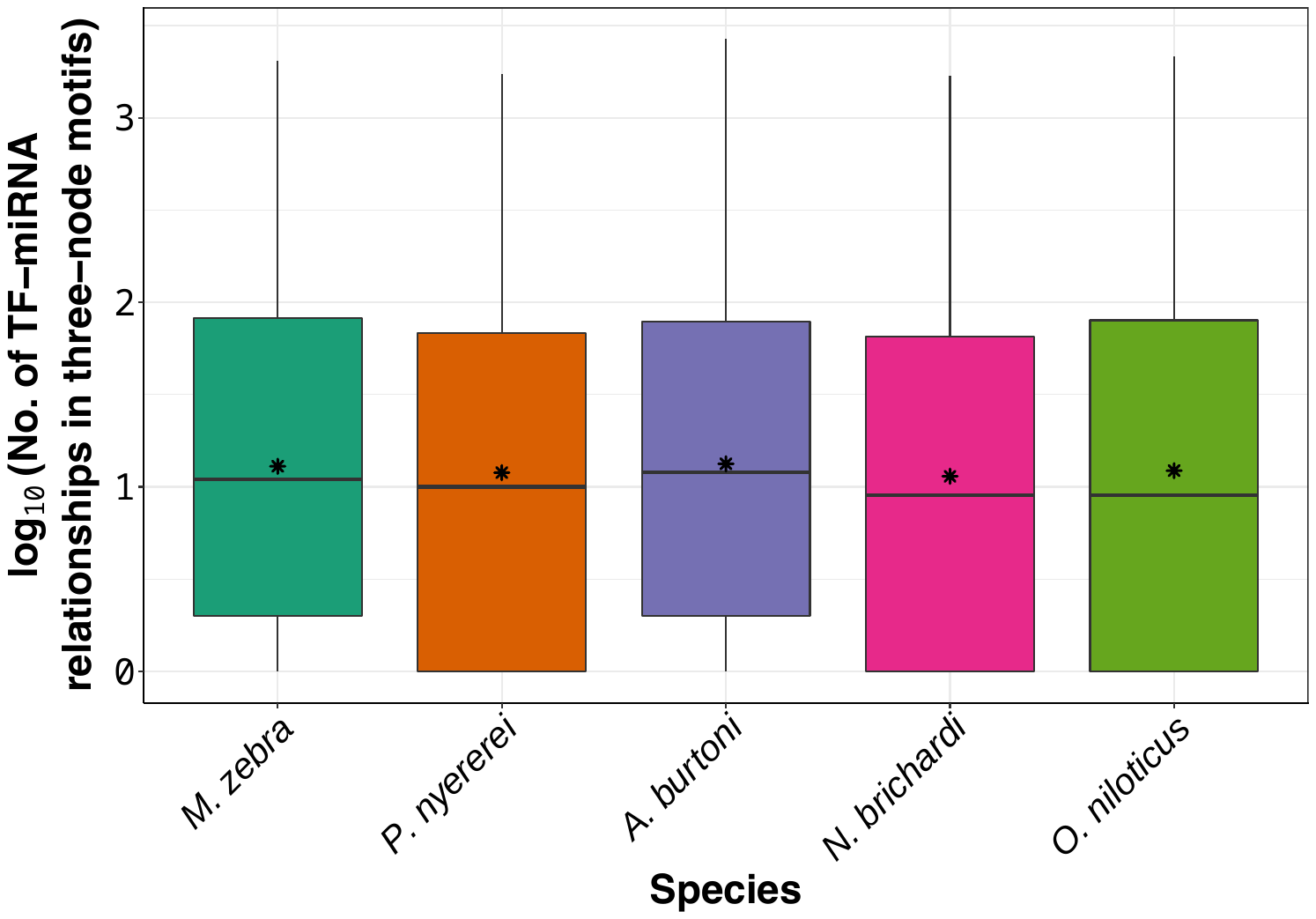

**Fig. S6 – Number of TF-miRNA relationships in three-node motifs of all genes in each species.** Boxplot of log_10_ counts of TF-miRNA relationships in three-node motifs (y-axis) in each species (x-axis) for 37,320,950 three-node motif edges (of TFs, TGs and miRNAs).

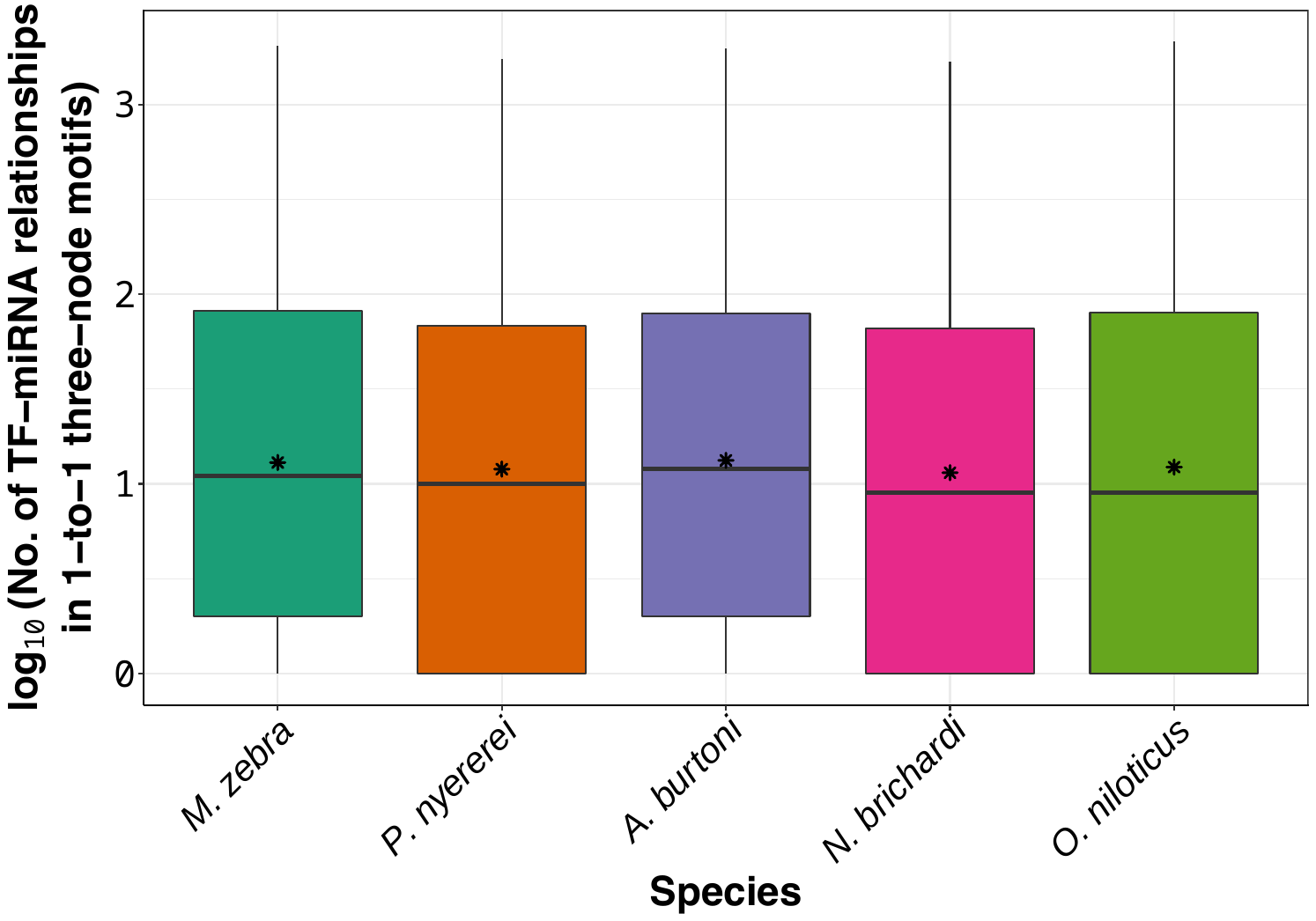

**Fig. S7 – Number of TF-miRNA relationships in three-node motifs of 1-to-1 orthologous genes in each species.** Boxplot of log_10_ counts of TF-miRNA relationships in three-node motifs (y-axis) in each species (x-axis) for 17,987,294 three-node motif edges composed of 1-to-1 orthologous edges (of TFs, TGs and miRNAs).

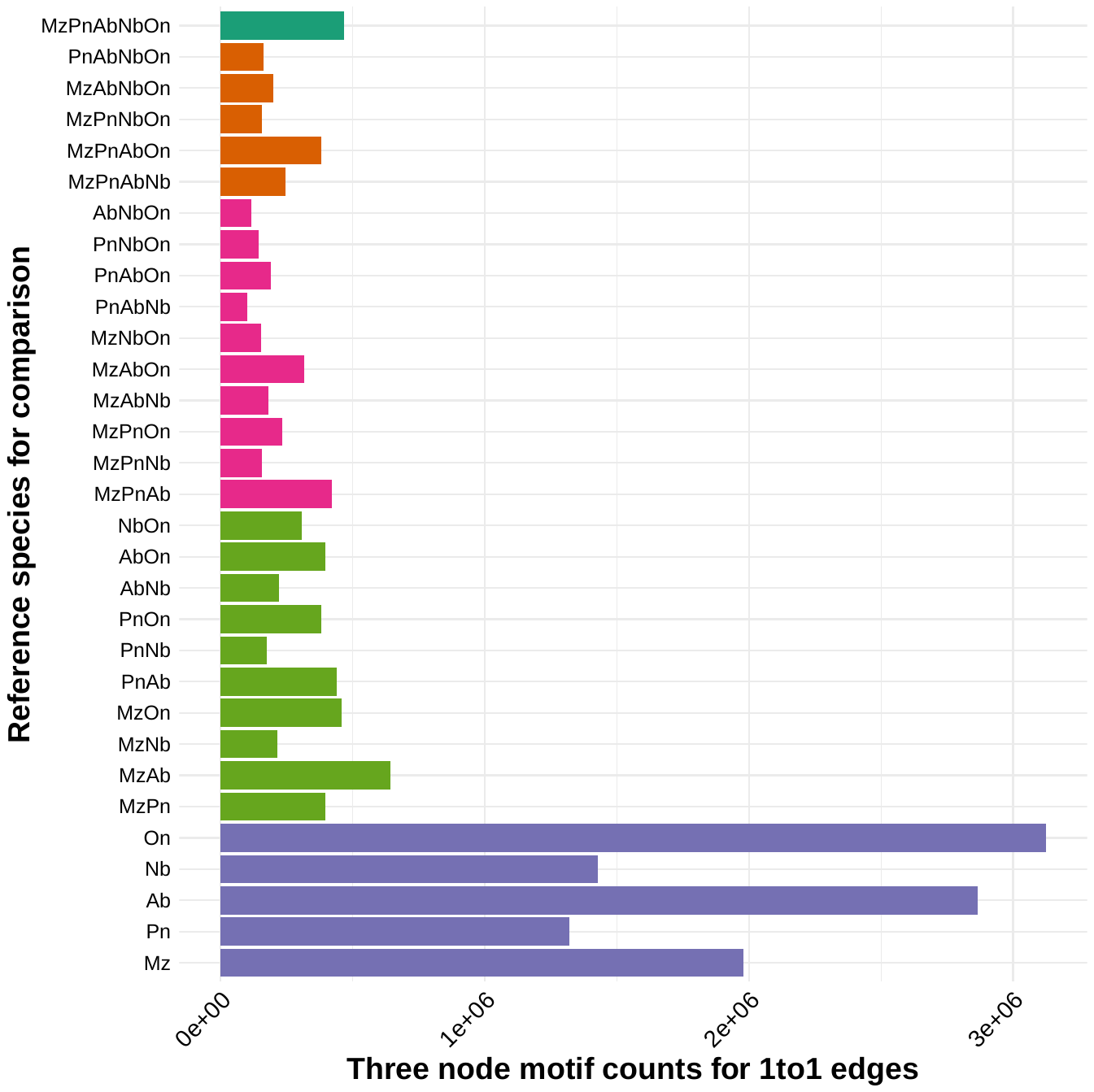

**Fig. S8 – Conserved and novel motifs between five species comparisons for 1-to-1 orthologous node edges.** Bar plot using 1-to-1 orthologous node edges shows counts of motifs in single species (purple bars), two species (light green bars), three species (pink bars), four species (orange bars) and all five species (dark green bar) are shown. Species names have been abbreviated: On = *O. niloticus,* Nb = *N. brichardi*, Ab = *A. burtoni*, Pn = *P. nyererei* and Mz = *M. zebra.* All details included in Supplementary Table S2.

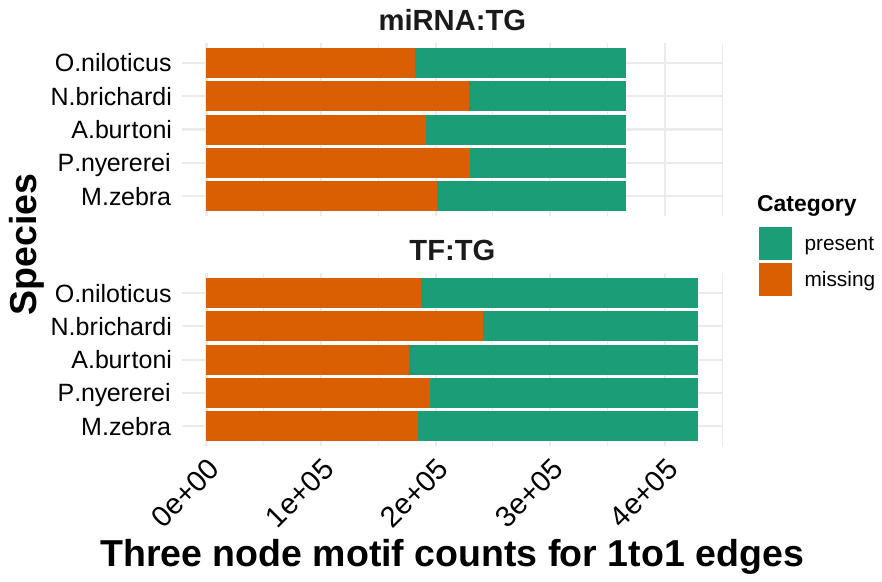

**Fig. S9 – Number of present and absent edges for each species 1-to-1 orthologous TG.** Bar plot using 1-to-1 orthologous node edges shows number of present and absent edges for each species 1-to-1 orthologous TG.

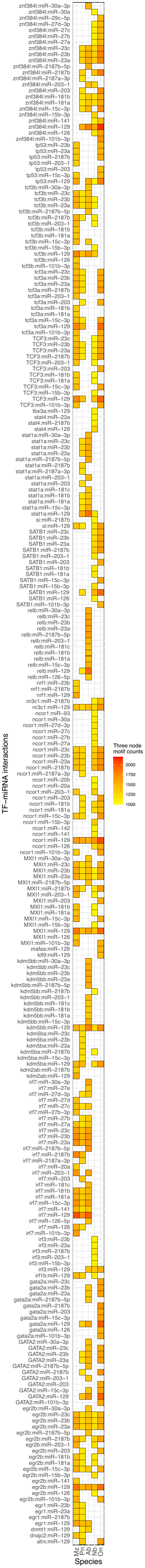
**Fig. S10 – Top 100 three-node motifs in species network edges.** Three-node motifs defined as transcription factor (TF) > target gene (TG) < miRNA relationships in species-specific edges of 1-to-1 orthologous nodes across all five species. Counts of TF-miRNA interactions (y-axis) in three-node motifs of all five species (x-axis). Grid colors indicate counts as per legend on *right.*

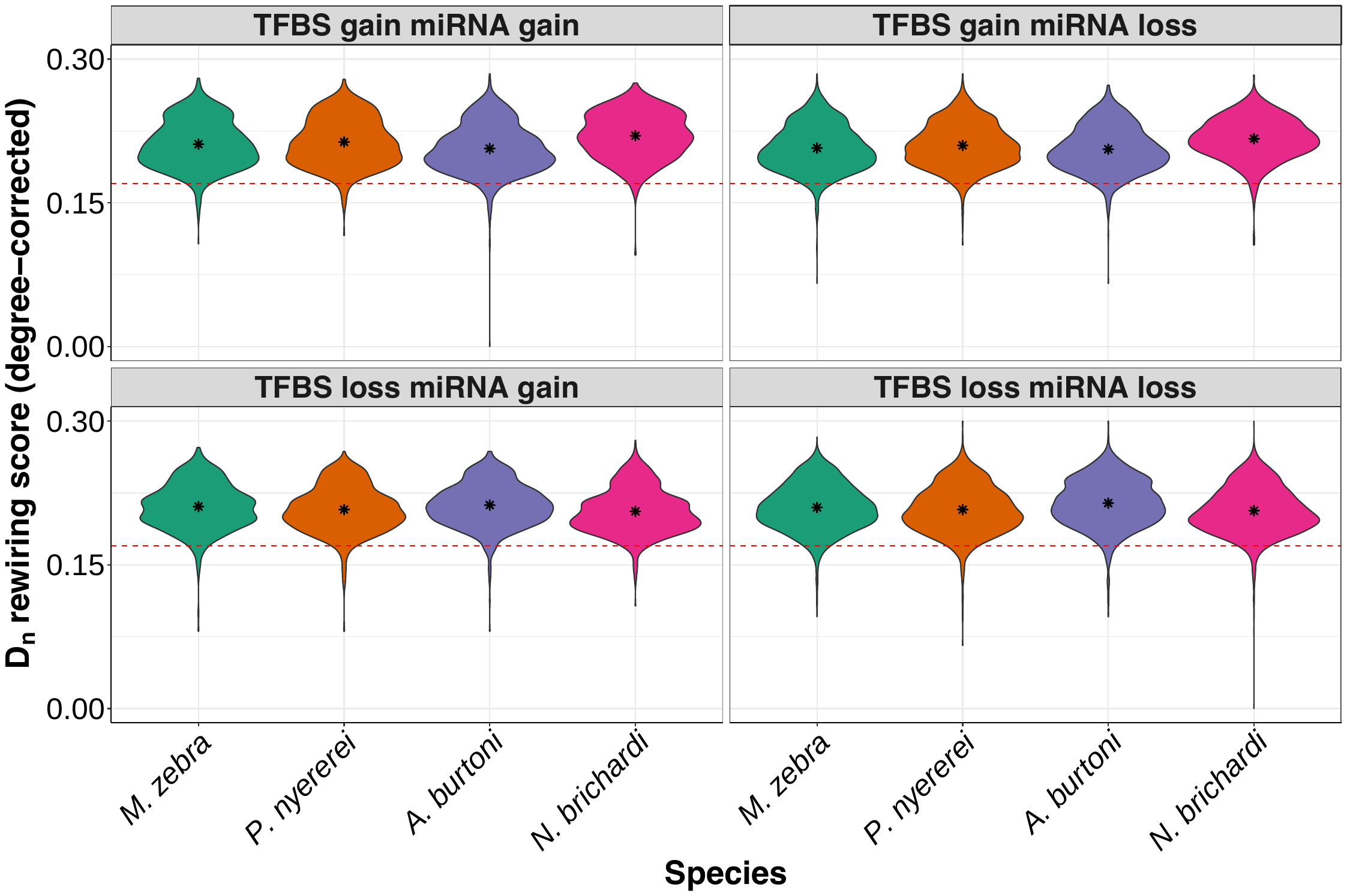

**Fig. S11 – Different models of TFBS and miRNA binding site evolution with associated rewiring rates of 1-to-1 orthogroups in four cichlids.** Violin plots of 4/8 models of binding site evolution in each species (x-axis) with DyNet rewiring score of each 1-to-1 orthogroup as degree corrected D_n_ score (y-axis). Red dotted line demarcates a *D_n_* score threshold of 0.17 (for rewired vs low to non-rewired genes), which was set based on the mean *D_n_* score for all orthogroups and used as a measure of significantly rewired genes based on our previous study (Mehta et al. 2021). All statistics are included in Supplementary Table S5.

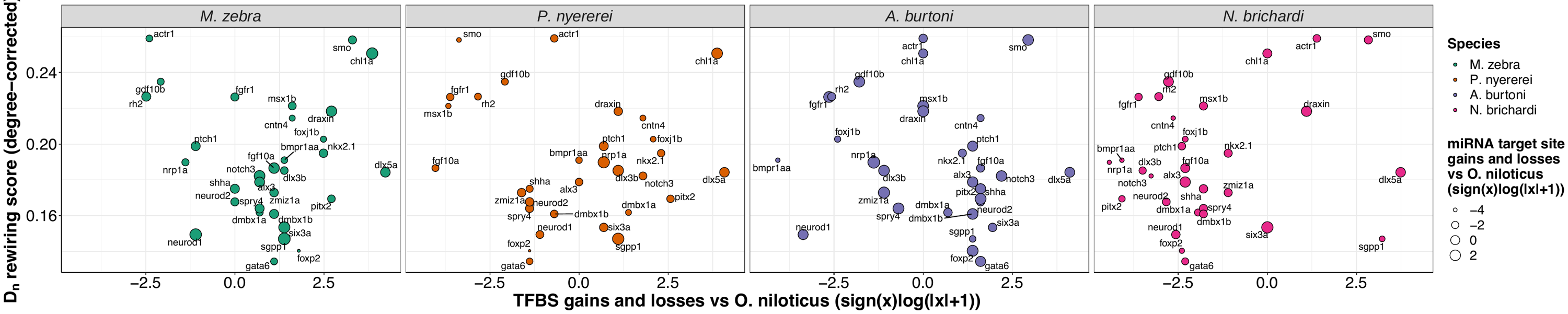

**Fig. S12 - Rewiring score and TFBS/miRNA binding site gain or loss in 1-to-1 orthologous candidate genes in four cichlids**. DyNet rewiring score as degree corrected D_n_ score (y-axis) against sign(x)(log(x+1)) no. of TFBSs gained/lost (x-axis) and miRNA gain/loss as dot size in 1-to-1 orthologous candidate genes.

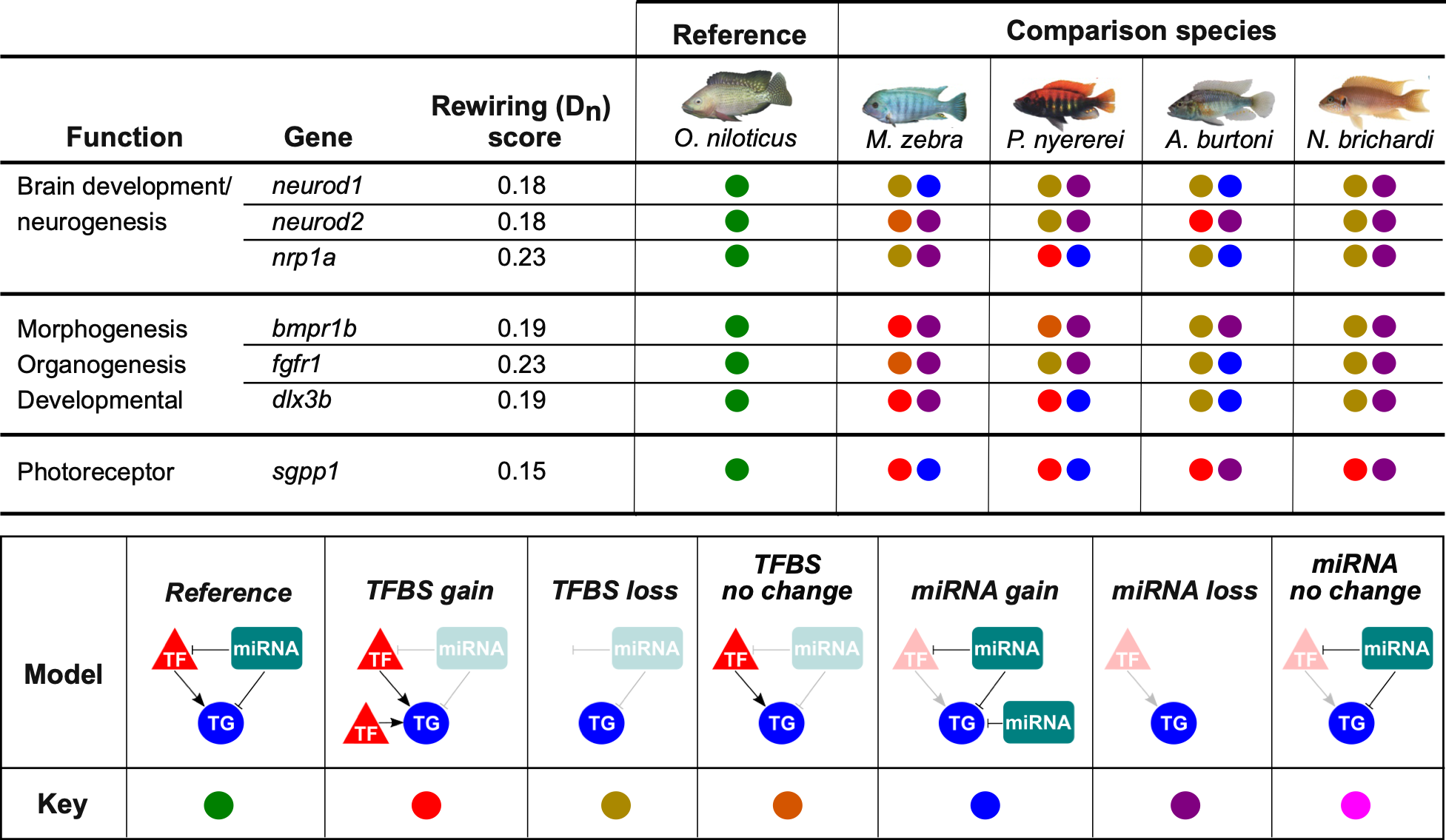

**Fig. S13 - Binding site evolution of seven cichlid adaptive trait genes.** DyNet rewiring (*D_n_*) score for all genes obtained from our previous study (Mehta et al. 2021). For the four comparison species, each genes model of TFBS and miRNA target site evolution in three-node motifs is calculated using the orthologous *O. niloticus* gene as a reference and demarcated as per the ‘model’ and ‘key’ in legend. All statistics are included in Supplementary Table S4-5.

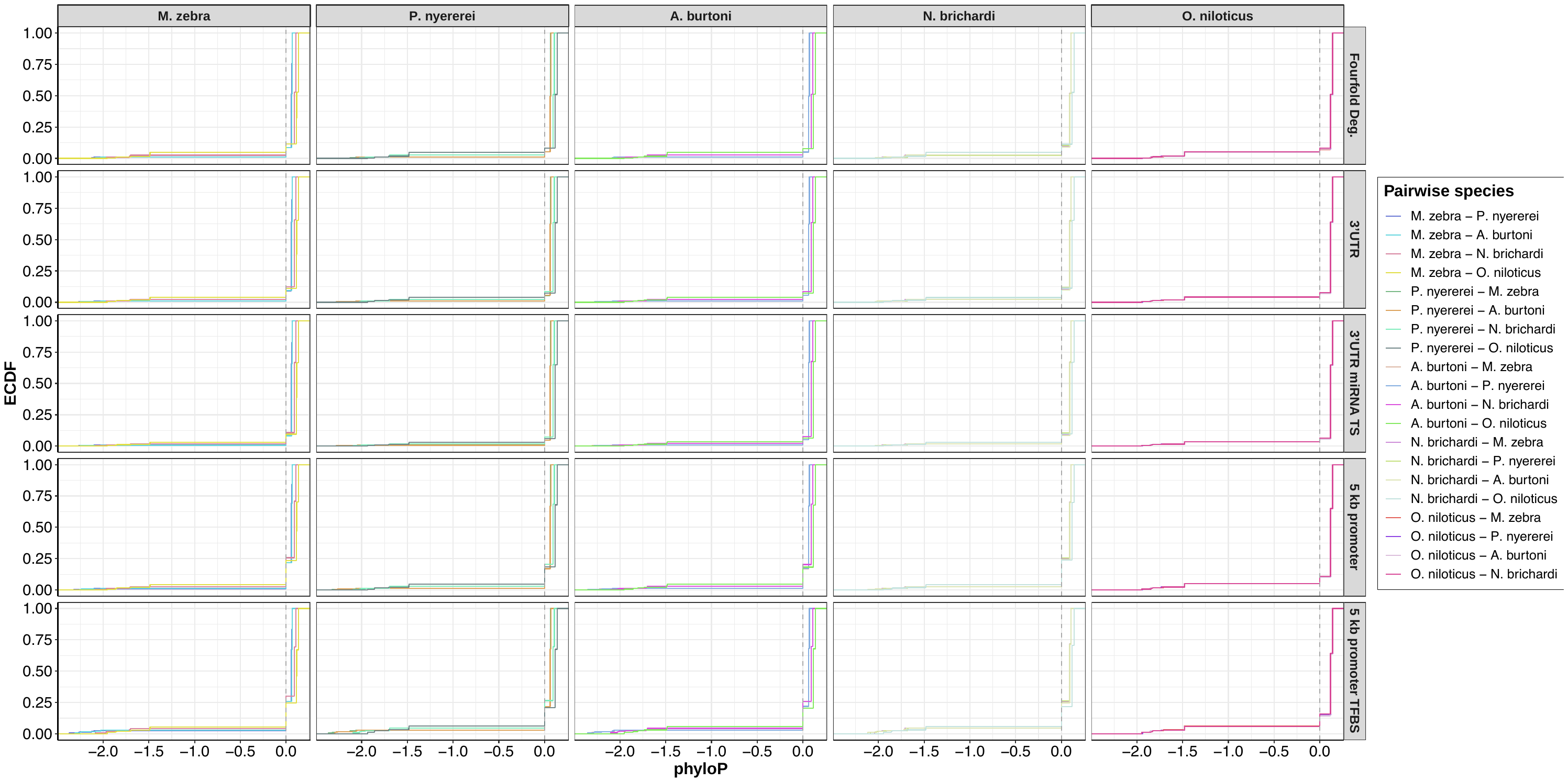

**Fig. S14 – Distribution of calculated conservation-acceleration (CONACC) scores using phyloP in coding and noncoding regulatory sites of the five cichlids.** Empirical cumulative distribution frequency (ECDF) in five features including fourfold degenerate sites and regulatory regions (3’ UTR, up to 5kb gene promoter, 3’ UTR miRNA target sites and 5kb gene promoter TFBSs) of distribution of frequency of CONACC scores across all five features in pairwise comparisons of all five species.

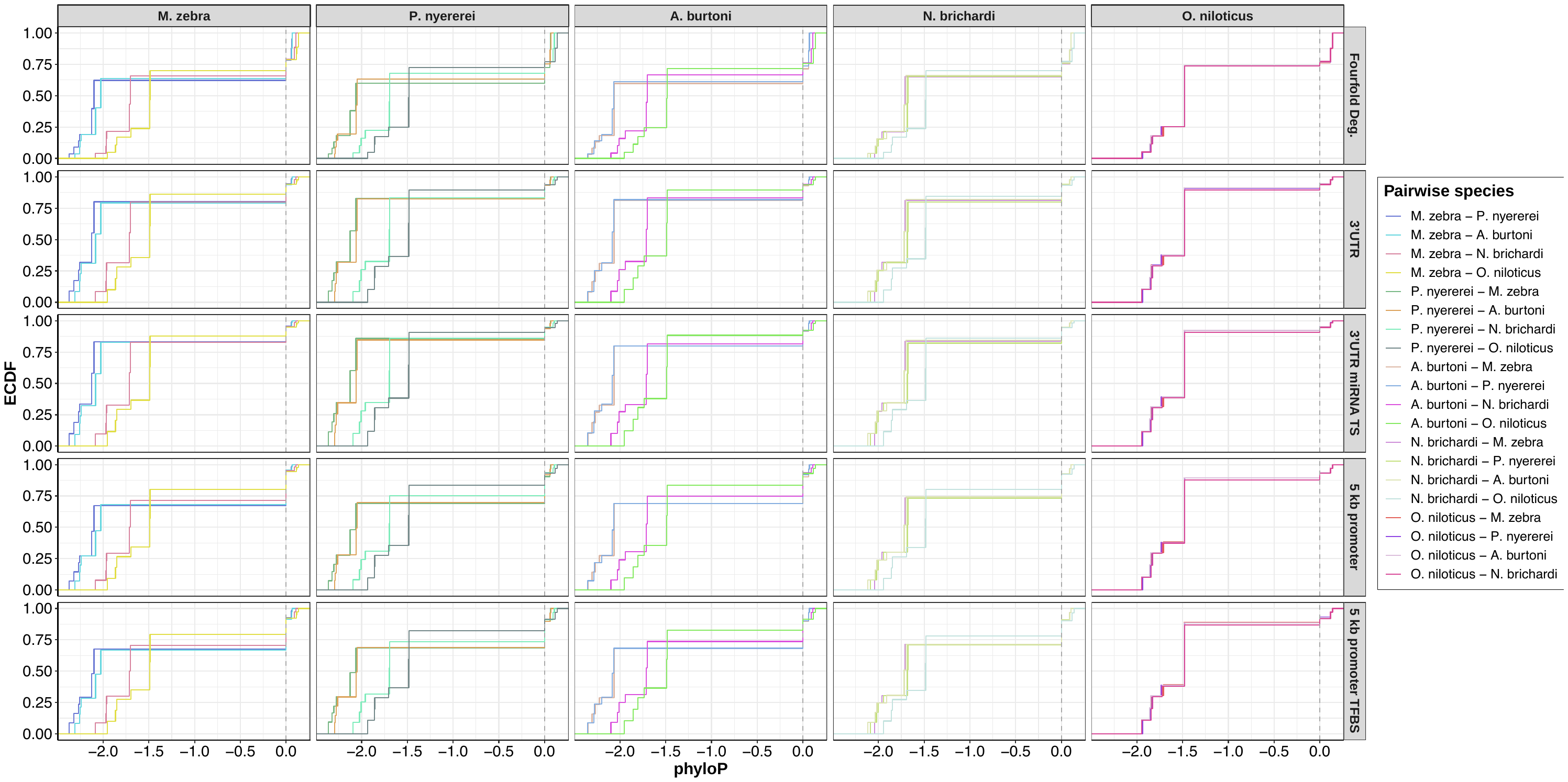

**Fig. S15 – *log d*istribution of calculated conservation-acceleration (CONACC) scores using phyloP in pairwise polymorphic sites overlapping coding and noncoding regulatory sites of the five cichlids.** Empirical cumulative distribution frequency (ECDF) in five features including fourfold degenerate sites and regulatory regions (3’ UTR, up to 5kb gene promoter, 3’ UTR miRNA target sites and 5kb gene promoter TFBSs) of distribution of frequency of CONACC scores of pairwise polymorphic sites in all five features in pairwise comparisons of all five species.

**Fig. S16** **- SNP genotypes overlapping ATF3 TFBS in *M. zebra sws1* promoter and other Lake Malawi species.** Lake Malawi phylogeny reproduced from published least controversial and all included species ASTRAL phylogeny (Malinsky et al. 2018), including *O. niloticus* as an outgroup. Phylogenetic branches labelled with species sample name and clade according to legends (*right*): A) Species foraging/diet habit (colour) (Hofmann et al. 2009) and phased SNP genotype (shape) (Malinsky et al. 2018); B) Adult opsin wavelength palette utilized (Hofmann et al. 2009); and C) species habitat (Hofmann et al. 2009; Froese and Pauly 2017). Ecological classifications are further described in Supplementary Table S19.

**Fig. S17** **- SNP genotypes overlapping miR-99a target site in *M. zebra sws1* 3’ UTR region and other Lake Malawi species.** Lake Malawi phylogeny reproduced from published least controversial and all included species ASTRAL phylogeny (Malinsky et al. 2018), including *P. nyererei, A. burtoni, N. brichardi* and *O. niloticus* as an outgroup. Phylogenetic branches labelled with species sample name and clade according to legends (*right*): A) Species foraging/diet habit (colour) (Hofmann et al. 2009) and phased SNP genotype (shape) (Malinsky et al. 2018); B) Adult opsin wavelength palette utilized (Hofmann et al. 2009); and C) species habitat (Hofmann et al. 2009; Froese and Pauly 2017). Ecological classifications are further described in Supplementary Table S19.

**Fig. S18** **- SNP genotypes overlapping MXI1 TFBS in *M. zebra rho* promoter and other Lake Malawi species.** Lake Malawi phylogeny reproduced from published least controversial and all included species ASTRAL phylogeny (Malinsky et al. 2018), including *P. nyererei* as an outgroup. Phylogenetic branches labelled with species sample name and clade according to legends (*right*): A) Species foraging/diet habit (colour) (Hofmann et al. 2009) and phased SNP genotype (shape) (Malinsky et al. 2018); and B) species habitat (Hofmann et al. 2009; Froese and Pauly 2017). Ecological classifications are further described in Supplementary Table S19.

**Fig. S19 - SNP genotypes overlapping miR-212 target site in *M. zebra rho* 3’ UTR region and other Lake Malawi species.** Lake Malawi phylogeny reproduced from published least controversial and all included species ASTRAL phylogeny (Malinsky et al. 2018), including *A. burtoni, N. brichardi* and *O. niloticus* as an outgroup. Phylogenetic branches labelled with species sample name and clade according to legends (*right*): A) Species foraging/diet habit (colour) (Hofmann et al. 2009) and phased SNP genotype (shape) (Malinsky et al. 2018); and B) species habitat (Hofmann et al. 2009; Froese and Pauly 2017). Ecological classifications are further described in Supplementary Table S19.

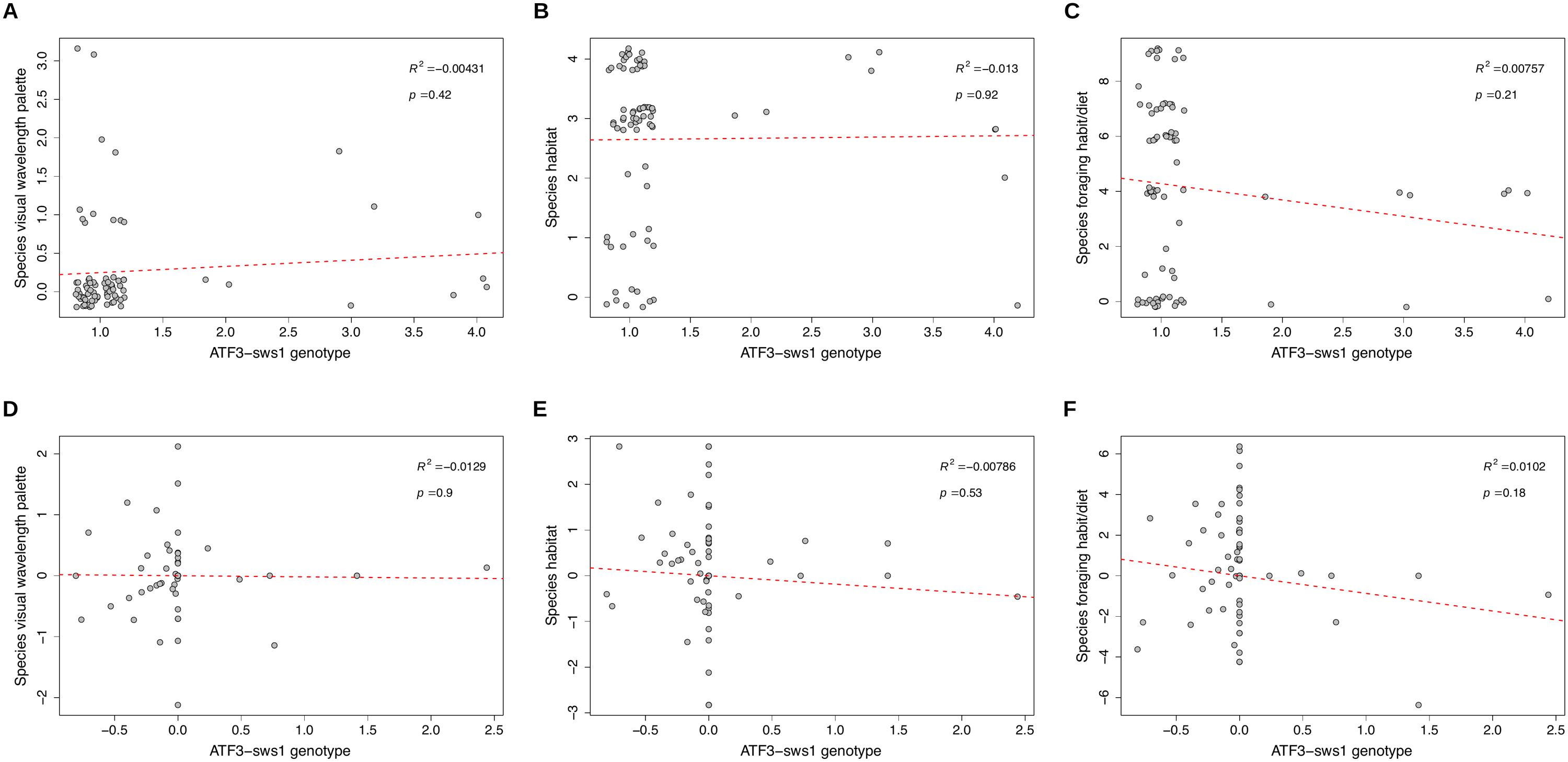

**Fig. S20 – Phylogenetic independent contrast analysis of ATF3-*sws1* TFBS genotypes of Lake Malawi species against their visual traits and ecology.** Phylogenetic independent scatterplots of ATF3-*sws1* TFBS genotypes (1=C|C, 2=C|T, 3=T|C, 4=T|T, 5=T/T) in 119 Lake Malawi individuals (73 species) against their respective **(a) visual wavelength palette** (0=N/A, 1=Short, 2=Medium, 3=Long); **(b) habitat** (0=N/A, 1=Rock, 2=Pelagic, 3=Benthopelagic, 4=Demersal); **(c) foraging habit/diet** (0=N/A, 1=Algae, 2=Aufwuchs, 3=Benthivore, 4=Fish, 5=Herbivore, 6=Invertebrates, 7=Mix, 8=Waste, 9=Zooplankton). Corresponding scatterplots of Lake Malawi ASTRAL phylogeny (Malinsky et al. 2018) and regression model fitted to ATF3-*sws1* TFBS genotypes (1=C|C, 2=C|T, 3=T|C, 4=T|T, 5=T/T) of 119 Lake Malawi individuals (73 species) against their respective **(d) visual wavelength palette** (0=N/A, 1=Short, 2=Medium, 3=Long); **(e) habitat** (0=N/A, 1=Rock, 2=Pelagic, 3=Benthopelagic, 4=Demersal); **(f) foraging habit/diet** (0=N/A, 1=Algae, 2=Aufwuchs, 3=Benthivore, 4=Fish, 5=Herbivore, 6=Invertebrates, 7=Mix, 8=Waste, 9=Zooplankton). All data points used as per Supplementary Fig. S16, with overlapping coordinates ‘jittered’ around their respective point to highlight density. Adjusted r^2^ and *p-*value of each regression line shown in top right of each plot. Ecological classifications are further described in Supplementary Table S19.

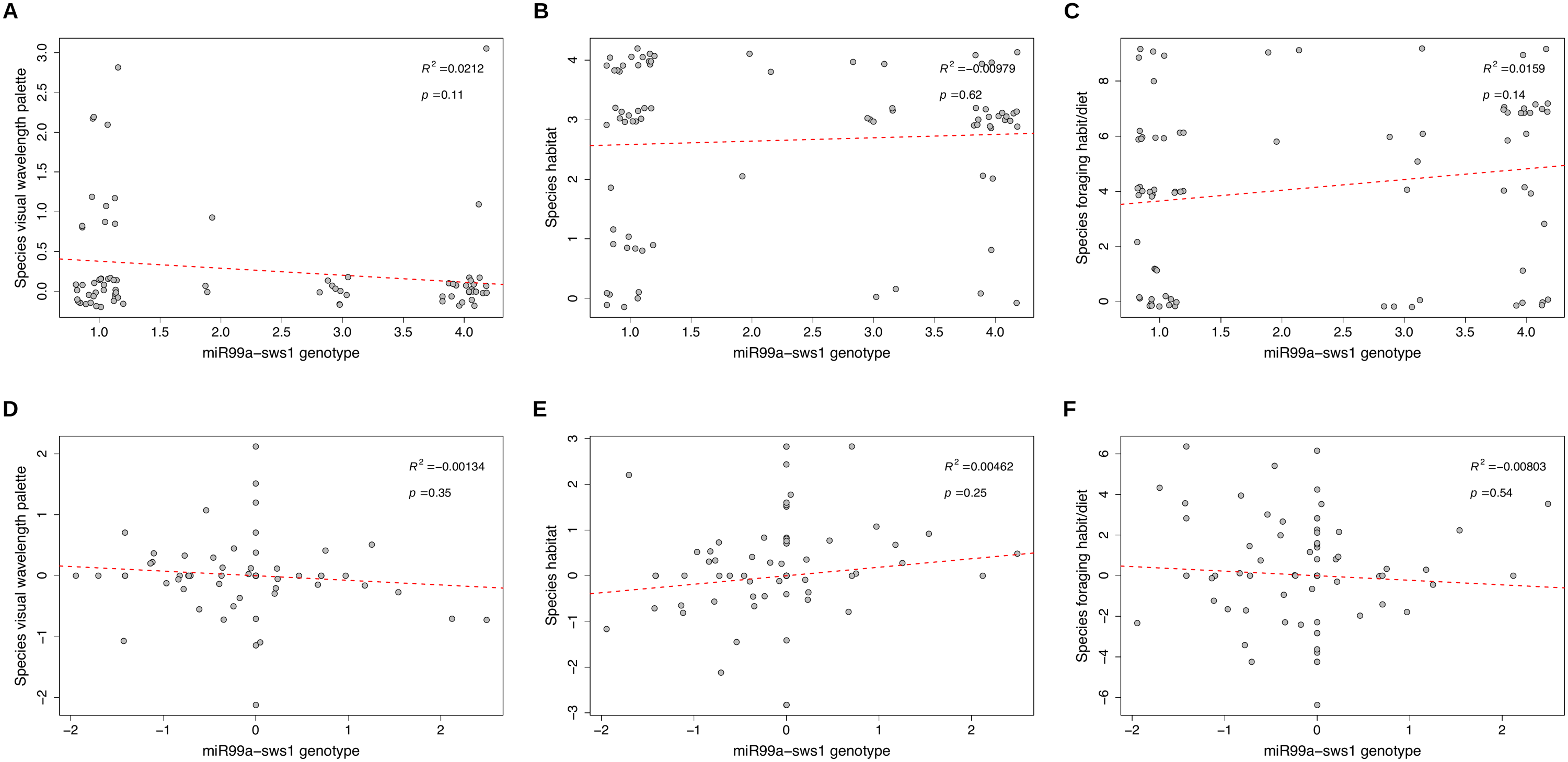

**Fig. S21 – Phylogenetic independent contrast analysis of miR-99a-*sws1* target site genotypes of Lake Malawi species against their visual traits and ecology.** Phylogenetic independent scatterplots of miR-99a-*sws1* target site genotypes (1=C|C, 2=C|A, 3=A|C, 4=A|A, 5=A/A) in 119 Lake Malawi individuals (73 species) against their respective **(a) visual wavelength palette** (0=N/A, 1=Short, 2=Medium, 3=Long); **(b) habitat** (0=N/A, 1=Rock, 2=Pelagic, 3=Benthopelagic, 4=Demersal); **(c) foraging habit/diet** (0=N/A, 1=Algae, 2=Aufwuchs, 3=Benthivore, 4=Fish, 5=Herbivore, 6=Invertebrates, 7=Mix, 8=Waste, 9=Zooplankton). Corresponding scatterplots of Lake Malawi ASTRAL phylogeny (Malinsky et al. 2018) and regression model fitted to miR-99a -*sws1* target site genotypes (1=C|C, 2=C|A, 3=A|C, 4=A|A, 5=A/A) of 119 Lake Malawi individuals (73 species) against their respective **(d) visual wavelength palette** (0=N/A, 1=Short, 2=Medium, 3=Long); **(e) habitat** (0=N/A, 1=Rock, 2=Pelagic, 3=Benthopelagic, 4=Demersal); **(f) foraging habit/diet** (0=N/A, 1=Algae, 2=Aufwuchs, 3=Benthivore, 4=Fish, 5=Herbivore, 6=Invertebrates, 7=Mix, 8=Waste, 9=Zooplankton). All data points used as per Supplementary Fig. S17, with overlapping coordinates ‘jittered’ around their respective point to highlight density. Adjusted r^2^ and *p-*value of each regression line shown in top right of each plot. Ecological classifications are further described in Supplementary Table S19.

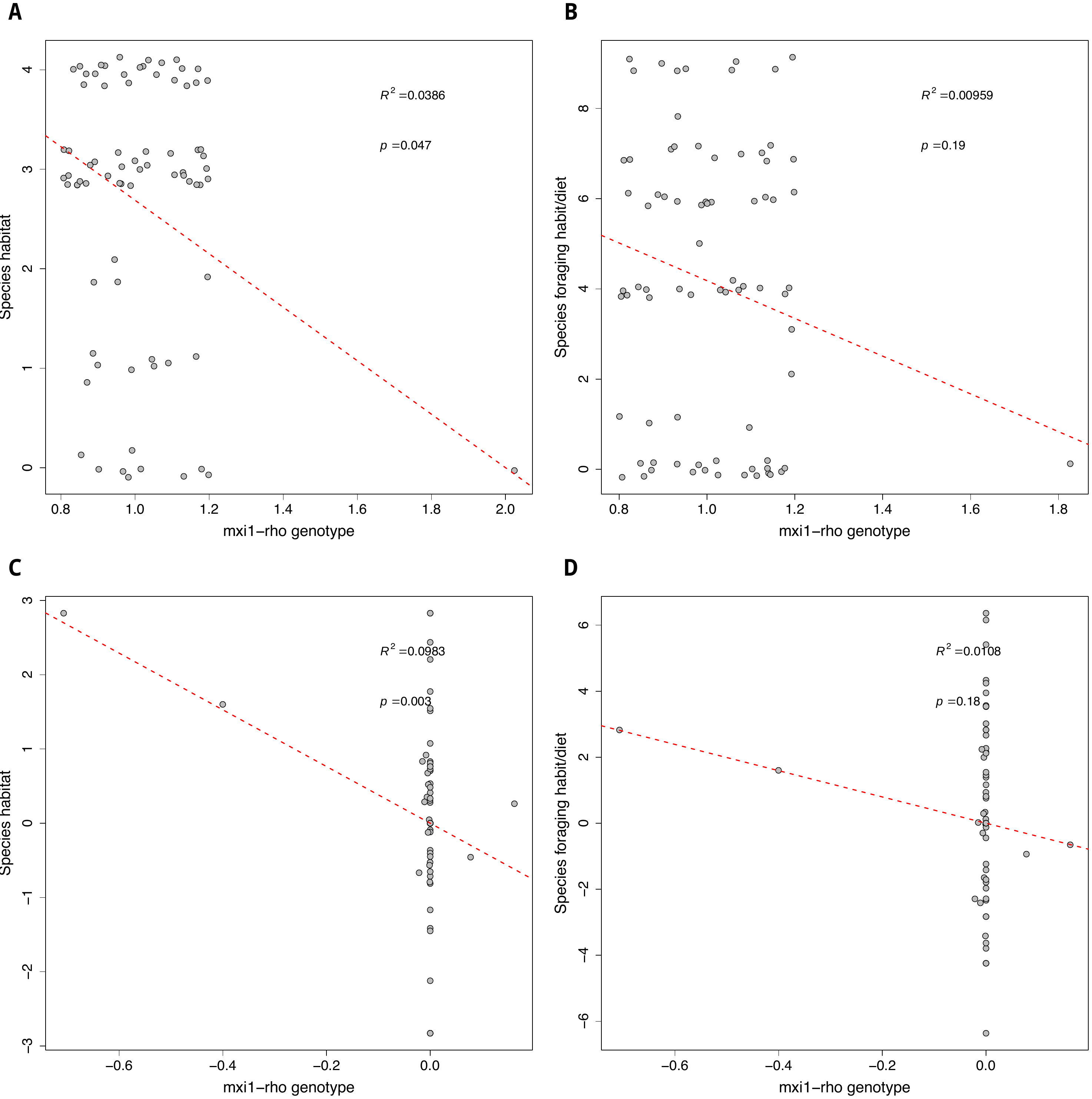

**Fig. S22 – Phylogenetic independent contrast analysis of MXI1-*rho* TFBS genotypes of Lake Malawi species against their visual traits and ecology.** Phylogenetic independent scatterplots of MXI1-*rho* target site genotypes (1=T|T, 2=C|T, 3=G/G) in 119 Lake Malawi individuals (73 species) against their respective **(a) habitat** (0=N/A, 1=Rock, 2=Pelagic, 3=Benthopelagic, 4=Demersal); **(b) foraging habit/diet** (0=N/A, 1=Algae, 2=Aufwuchs, 3=Benthivore, 4=Fish, 5=Herbivore, 6=Invertebrates, 7=Mix, 8=Waste, 9=Zooplankton). Corresponding scatterplots of Lake Malawi ASTRAL phylogeny (Malinsky et al. 2018) and regression model fitted to MXI1-*rho* target site genotypes (1=T|T, 2=C|T, 3=G/G) of 119 Lake Malawi individuals (73 species) against their respective **(c) habitat** (0=N/A, 1=Rock, 2=Pelagic, 3=Benthopelagic, 4=Demersal); **(d) foraging habit/diet** (0=N/A, 1=Algae, 2=Aufwuchs, 3=Benthivore, 4=Fish, 5=Herbivore, 6=Invertebrates, 7=Mix, 8=Waste, 9=Zooplankton). All data points used as per Supplementary Fig. S18, with overlapping coordinates ‘jittered’ around their respective point to highlight density. Adjusted r^2^ and *p-*value of each regression line shown in top right of each plot. Ecological classifications are further described in Supplementary Table S19.

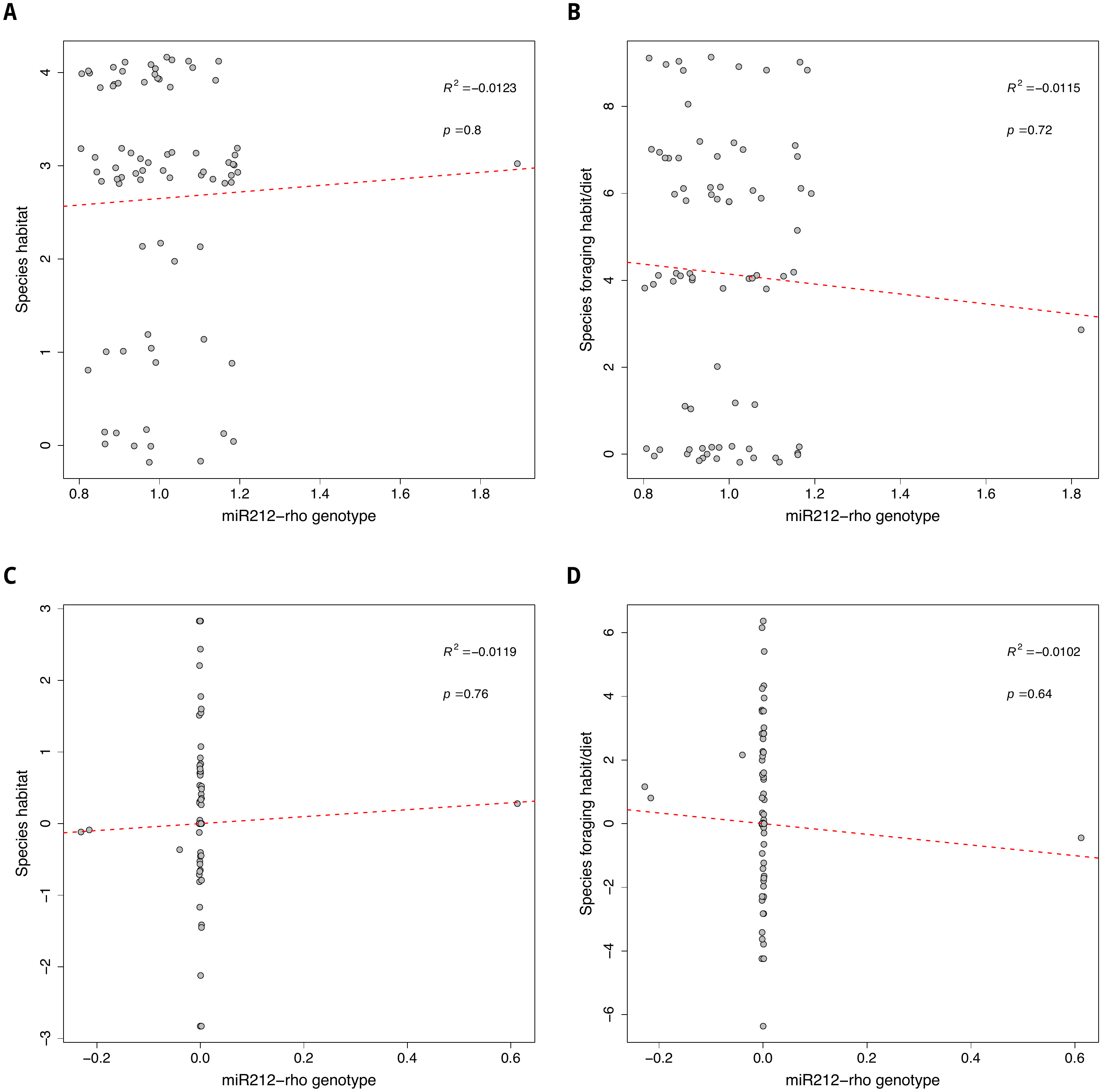

**Fig. S23 – Phylogenetic independent contrast analysis of miR-212-*rho* target site genotypes of Lake Malawi species against their visual traits and ecology.** Phylogenetic independent scatterplots of miR-212-*rho* target site genotypes (1=A|A, 2=C|C, 3=C/C) in 119 Lake Malawi individuals (73 species) against their respective **(a) habitat** (0=N/A, 1=Rock, 2=Pelagic, 3=Benthopelagic, 4=Demersal); **(b) foraging habit/diet** (0=N/A, 1=Algae, 2=Aufwuchs, 3=Benthivore, 4=Fish, 5=Herbivore, 6=Invertebrates, 7=Mix, 8=Waste, 9=Zooplankton). Corresponding scatterplots of Lake Malawi ASTRAL phylogeny (Malinsky et al. 2018) and regression model fitted to miR-212-*rho* target site genotypes (1=A|A, 2=C|C, 3=C/C) of 119 Lake Malawi individuals (73 species) against their respective **(c) habitat** (0=N/A, 1=Rock, 2=Pelagic, 3=Benthopelagic, 4=Demersal); **(d) foraging habit/diet** (0=N/A, 1=Algae, 2=Aufwuchs, 3=Benthivore, 4=Fish, 5=Herbivore, 6=Invertebrates, 7=Mix, 8=Waste, 9=Zooplankton). All data points used as per Supplementary Fig. S19, with overlapping coordinates ‘jittered’ around their respective point to highlight density. Adjusted r^2^ and *p-*value of each regression line shown in top right of each plot. Ecological classifications are further described in Supplementary Table S19.

**Fig. S24 - SNP genotypes overlapping DNAJC2 TFBS in *M. zebra msx1b* promoter and other Lake Malawi species.** Lake Malawi phylogeny reproduced from published least controversial and all included species ASTRAL phylogeny (Malinsky et al. 2018), including *N. brichardi* as an outgroup. Phylogenetic branches labelled with species sample name and clade according to legends (*right*): A) Species foraging/diet habit (colour) (Hofmann et al. 2009) and phased SNP genotype (shape) (Malinsky et al. 2018); and B) species habitat (Hofmann et al. 2009; Froese and Pauly 2017). Ecological classifications are further described in Supplementary Table S19.

**Fig. S25 - SNP genotypes overlapping miR-129 target site in *M. zebra msx1b* 3’ UTR region and other Lake Malawi species.** Lake Malawi phylogeny reproduced from published least controversial and all included species ASTRAL phylogeny (Malinsky et al. 2018), including *P. nyererei, A. burtoni* and *N. brichardi* as an outgroup. Phylogenetic branches labelled with species sample name and clade according to legends (*right*): A) Species foraging/diet habit (colour) (Hofmann et al. 2009) and phased SNP genotype (shape) (Malinsky et al. 2018); and B) species habitat (Hofmann et al. 2009; Froese and Pauly 2017). Ecological classifications are further described in Supplementary Table S19.

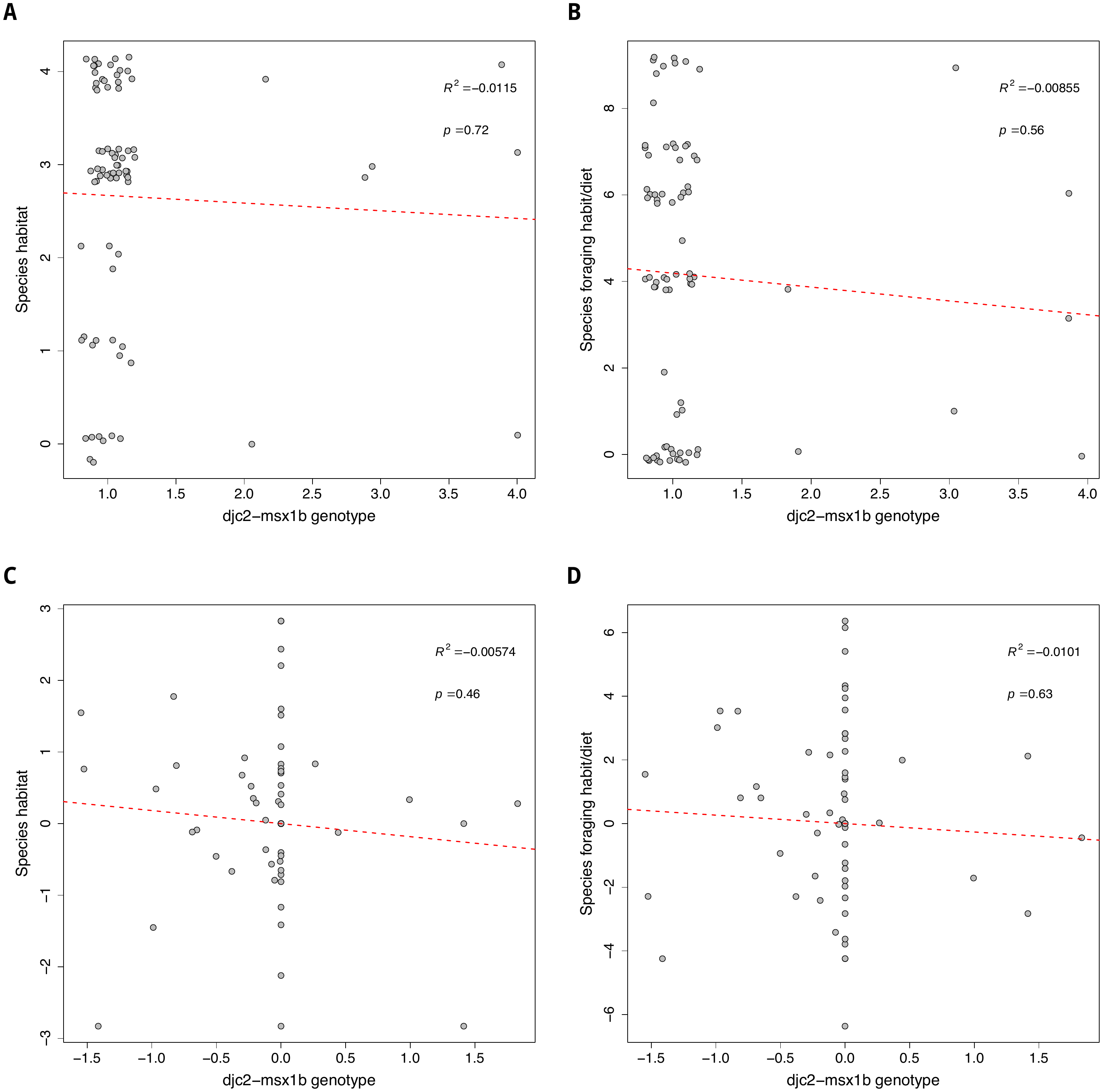

**Fig. S26 – Phylogenetic independent contrast analysis of DJC2-*msx1b* TFBS genotypes of Lake Malawi species against their visual traits and ecology.** Phylogenetic independent scatterplots of DJC2-*msx1b* TFBS genotypes (1=G|G, 2=T|G, 3=G|T, 4=T|T) in 119 Lake Malawi individuals (73 species) against their respective **(a) habitat** (0=N/A, 1=Rock, 2=Pelagic, 3=Benthopelagic, 4=Demersal); **(b) foraging habit/diet** (0=N/A, 1=Algae, 2=Aufwuchs, 3=Benthivore, 4=Fish, 5=Herbivore, 6=Invertebrates, 7=Mix, 8=Waste, 9=Zooplankton). Corresponding scatterplots of Lake Malawi ASTRAL phylogeny (Malinsky et al. 2018) and regression model fitted to DJC2-*msx1b* TFBS genotypes (1=G|G, 2=T|G, 3=G|T, 4=T|T) of 119 Lake Malawi individuals (73 species) against their respective **(c) habitat** (0=N/A, 1=Rock, 2=Pelagic, 3=Benthopelagic, 4=Demersal); **(d) foraging habit/diet** (0=N/A, 1=Algae, 2=Aufwuchs, 3=Benthivore, 4=Fish, 5=Herbivore, 6=Invertebrates, 7=Mix, 8=Waste, 9=Zooplankton). All data points used as per Supplementary Fig. S24, with overlapping coordinates ‘jittered’ around their respective point to highlight density. Adjusted r^2^ and *p-*value of each regression line shown in top right of each plot. Ecological classifications are further described in Supplementary Table S19.

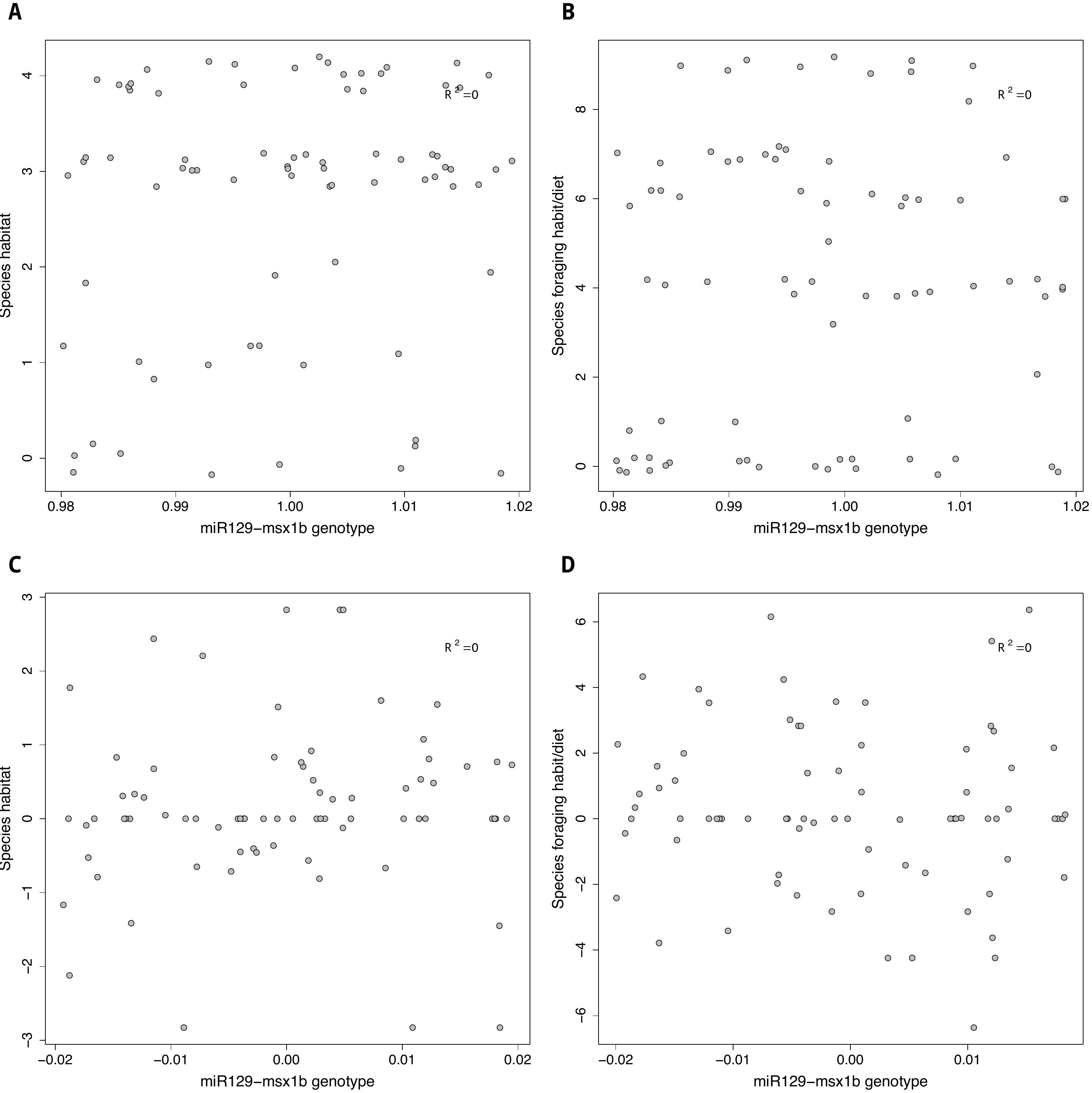

**Fig. S27 – Phylogenetic independent contrast analysis of miR-129-*msx1b* target site genotypes of Lake Malawi species against their visual traits and ecology.** Phylogenetic independent scatterplots of miR-129-*msx1b* target site genotypes (1=A|A, 2=A|C, 3=C/C) in 119 Lake Malawi individuals (73 species) against their respective **(a) habitat** (0=N/A, 1=Rock, 2=Pelagic, 3=Benthopelagic, 4=Demersal); **(b) foraging habit/diet** (0=N/A, 1=Algae, 2=Aufwuchs, 3=Benthivore, 4=Fish, 5=Herbivore, 6=Invertebrates, 7=Mix, 8=Waste, 9=Zooplankton). Corresponding scatterplots of Lake Malawi ASTRAL phylogeny (Malinsky et al. 2018) and regression model fitted to miR-129-*msx1b* target site genotypes (1=A|A, 2=A|C, 3=C/C) of 119 Lake Malawi individuals (73 species) against their respective **(c) habitat** (0=N/A, 1=Rock, 2=Pelagic, 3=Benthopelagic, 4=Demersal); **(d) foraging habit/diet** (0=N/A, 1=Algae, 2=Aufwuchs, 3=Benthivore, 4=Fish, 5=Herbivore, 6=Invertebrates, 7=Mix, 8=Waste, 9=Zooplankton). All data points used as per Supplementary Fig. S25, with overlapping coordinates ‘jittered’ around their respective point to highlight density. Since genotypes for species used are the same, there is no regression line for these plots. Ecological classifications are further described in Supplementary Table S19.
